# A quantitative description for optical mass measurement of single biomolecules

**DOI:** 10.1101/2023.03.28.534430

**Authors:** Jan Becker, Jack S. Peters, Ivor Crooks, Seham Helmi, Marie Synakewicz, Benjamin Schuler, Philipp Kukura

## Abstract

Label-free detection of single biomolecules in solution has been achieved using a variety of experimental approaches over the past decade. Yet, our understanding of the magnitude of the optical contrast and its relationship to the underlying atomic structure, as well as the achievable measurement sensitivity and precision remain poorly defined. Here, we use a Fourier optics approach combined with an atomic structure-based molecular polarizability model to simulate mass photometry experiments from first principles. We find excellent agreement between several key experimentally-determined parameters such as optical contrast-to-mass conversion, achievable mass accuracy and molecular shape/orientation dependence. This allows us to determine detection sensitivity and measurement precision that is mostly independent of the optical detection approach chosen, resulting in a general framework for light-based single molecule detection and quantification.

## Introduction

Recent developments in ultrasensitive light microscopy^1,2^ have enabled the quantification of biomolecular mass, charge and size at the single molecule level and in solution^3–8^. In mass photometry (MP) light scattered from a protein when it binds to or moves along a glass coverslip in solution is detected together with partially reflected light from the glass-water interface (**Fig. 1a**). MP has demonstrated both high mass accuracy and precision on the order of a few percent of the object mass, enabled by high measurement precision at the single molecule level^3^. This has been achieved through the selective attenuation of the reflected light using a mask in the back-focal-plane (BFP) of the optical system^9^, coupled with averaging of detected photoelectrons by the imaging camera and post-processing of the raw images, enabling the detection and resolution of oligomeric states and protein complexes^10^. As a result, MP can be used to quantify interaction affinities and kinetics,^11^ molecular organisation^12^, and for studies of biomolecular dynamics^13^.

**Figure 1:**
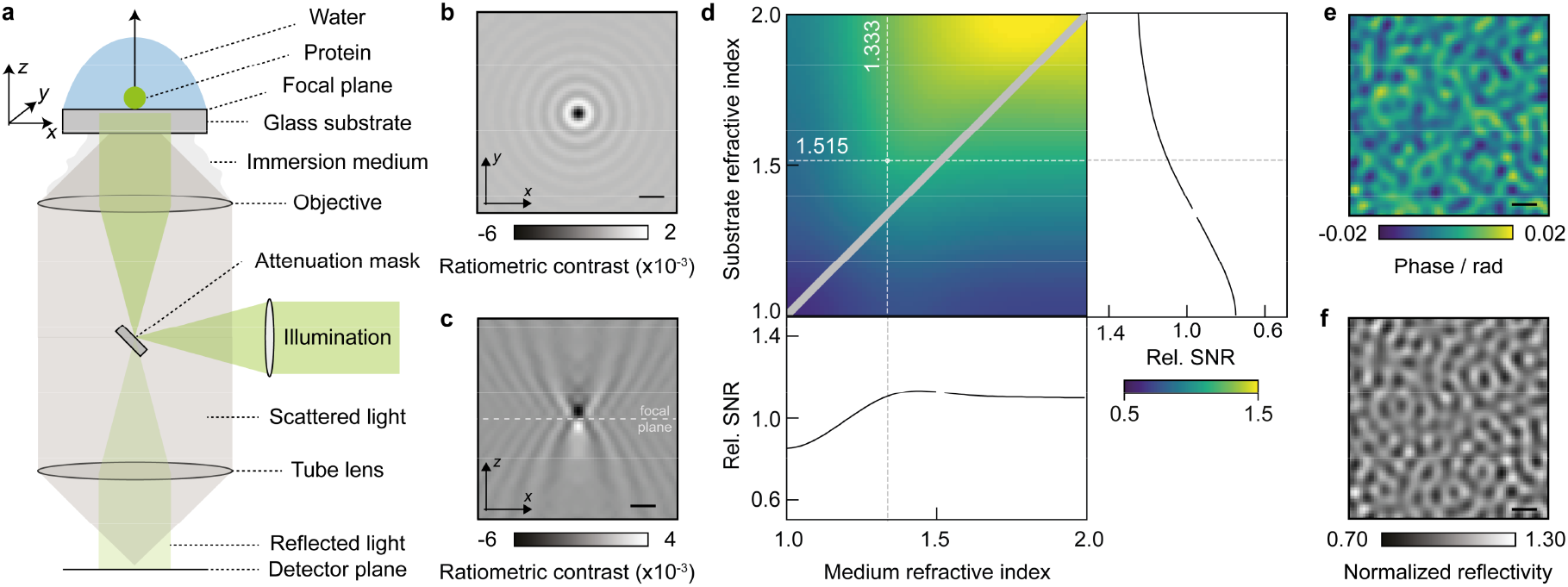
Fundamentals of image formation in a contrast-enhanced back reflection geometry. **a** Schematic representation of the simulated, widefield, mass photometry (MP) setup, including an attenuation mask in the back-focal-plane (BFP) of the objective lens, which selectively reduces light reflected from the glass coverslip. **b,c** Simulated ratiometric contrast for a single 24-mer of the small heat shock protein HSP16.5 (m = 396 kDa) at an atomically flat glass-water interface. **d** Relative signal-to-noise ratio (SNR) when varying the refractive indices of substrate and buffer medium. The diagonal indicates refractive index matching, where no reference field is available due to a lack of reflection. **e** Simulated phase retardation map at λ = 445 nm arising from nanoscopic roughness of glass coverslips on the order of ∼ 2 nm height variations over ∼ 100 nm length scales. **f** Resulting speckle-like image using a 0.1% transmission mask, normalized to the expected reflectivity from a flat glass-water interface (see Supplement 6). Scalebars = 0.5 μm.

The molecular mass, *m*, for unknown samples is inferred from the optical contrast of the molecule under investigation and an empirical scaling between the contrast and species of known molecular mass. This relationship can be approximated by estimating the excess polarizability *α* using the refractive index of proteins *n*_*p*_ and assuming a spherical shape^14^. Such a description, however, does not consider the exact shape and atomic structure of the protein, the influence of an interface on the scattering properties of the object, nor the details of the optical arrangement used to perform the measurement. In addition, the refractive index of a single protein is poorly defined. As a result, to date there is no quantitative, molecular-level description of light-based mass measurement and what properties define the limits and opportunities in measurement sensitivity, precision, and accuracy.

We thus set out to develop an approach capable of simulating images of individual proteins on microscope cover glass in an MP instrument coupled with an explicit atomic description of the molecular polarizability, and thereby explore key aspects such as: 1. To which degree optical contrasts reported to date experimentally match those predicted by theory. 2. How the measured signal depends on molecular shape & orientation, thereby informing on the ultimately achievable mass accuracy and resolution. 3. The current and likely future limits on measurement sensitivity and resolution for light-based single molecule characterisation for MP and beyond.

## Results

Our model simulates an experimental setup based on plane wave illumination, a (simplified) single lens high numerical aperture (*NA*) objective for light delivery and collection that is refractive index matched to the sample interface (here glass; *n*_*g*_), while the protein under investigation is embedded in a medium of refractive index *n*_*m*_ (e.g. water). A spatial mask is used to selectively attenuate light reflected from the coverslip substrate, which increases the optical contrast, simplifies the accurate determination of the focal position for maximum contrast, and allows for higher illumination powers given limited camera full well capacities (**Fig. 1a**)^11^. We mathematically model the influence of the imaging system as the following convolution operation (defined as: 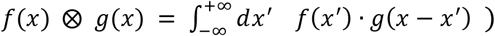 with an amplitude point-spread-function 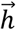 (APSF)^15^ :

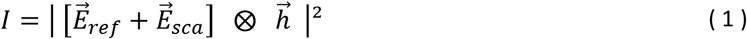

yielding the detectable intensity, *Ī* with 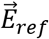 and 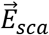 being the reflected and scattered electric fields, respectively. These are directly linked to the illuminating field 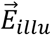 through:

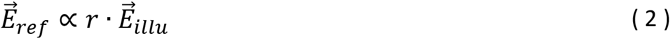

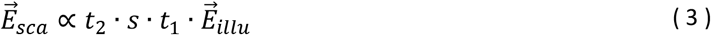

where *r* and *t* are the Fresnel coefficients for reflection and transmission^16^ and the subscripts indicate transmission of 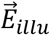 from glass → water (*t*_1_) or that of the scattered field 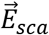 from water → glass (*t*_2_). Note that the influence of the attenuation mask is realized by propagating both, 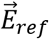 and 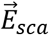, into the BFP of the objective lens, where they are being multiplied by a circular mask (corresponding to an effective numerical aperture, *NA*, of 0.58) with a given transmission strength *τ* ^2^ (1%; unless otherwise stated), motivated by experimental parameters^9^. The scattering coefficient is given by the polarizability *α* of the protein, which in the Rayleigh regime^14^ can be approximated to be proportional to the particle volume:

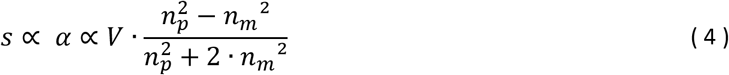

Note that our description of image formation mainly differs from experimental MP setups through non-scanned illumination, which is experimentally used to allow for widefield illumination without being limited by speckle-artifacts, and minimise the spatial extent of the molecular point-spread function^3^. Further details on the employed theoretical model, such as including the near-field effects of the protein scattering at a refractive index interface, high NA focusing effects and phase aberrations due to imaging into a layer of different refractive index (buffer vs glass) are given in the Methods 1 and the Supplementary Information 1 -8.

We begin our numerical investigation with Hsp16.5, a highly symmetric small heat shock protein, which forms spherical 24-mers of *m* = 396 kDa adsorbed on a glass substrate immersed in water to enable direct comparison with early experiments^9^. In a first iteration, we approximated the protein as a sphere of radius 5.6 nm and refractive index *n*_*p*_ = 1.480^17^. The substrate refractive index was chosen to be *n*_*g*_ = 1.515 as typical for borosilicate microscope cover glass used in MP^16^, but assumed to be atomically flat for simplicity. Experimentally, the limited full well capacity of CMOS imaging sensors requires both spatial and temporal summing of detected photoelectrons to reduce shot noise-induced background fluctuations and thereby optimise the attainable signal-to-noise ratio (SNR), taken as the ratio between the maximum signal amplitude introduced by the protein and the unknown background variations. In our simulations, we can take advantage of in principle unlimited full well capacities of the (virtual) camera pixels, which simplifies the image generation. The original experiments on Hsp16.5 used 2 × 2 spatial binning, corresponding to 60 nm/pixel at the reported magnification^9^, followed by summing photoelectrons from 100 subsequent images, leading to a total of 10^7^ detected photoelectrons per spatio-temporally binned camera pixel.

The quantity typically used to report the optical contrast is that of the *ratiometric* contrast *C*:

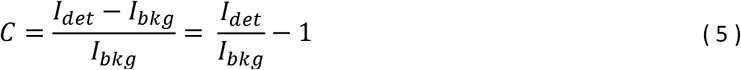

which is the relative difference between the measured intensity with (*I*_*det*_) and without (*I*_*bkg*_) the scatterer^9^. At the experimental illumination wavelength of *λ* = 445 nm, the simulated (here without shot noise; details on simulation parameters are given in Methods 2) ratiometric contrast of *C* ∼ 0.62% is close to the experimental result of 0.6%, as is the appearance of Airy rings arising from plane wave illumination (**Fig. 1b**). Note that the maximum contrast does not coincide with the nominal focus position (**Fig. 1c**), instead requiring a displacement of the sample by ∼190 nm along the optical axis to optimise the phase difference between scattered and reflected light by tuning the Gouy phase^18^. These results suggest that our model produces image contrasts in good agreement with experimental results, where the contrast is optimised by maximizing the (spatial) standard deviation of the glass roughness.

To explore the dependence of the image contrast on the refractive index of both the medium and the substrate, which in principle are tuneable away from that of water (*n*_*m*_ = 1.333) and borosilicate glass (*n*_*g*_ = 1.515), we varied both parameters and evaluated the achievable SNR for a constant power density incident on the sample. While increasing both refractive indices for a fixed refractive index of the protein (*n*_*p*_ = 1.46), which we assume to be non-tuneable, leads to a modest increase in the achievable SNR, the overall achievable increase is modest (**Fig. 1d**). Nevertheless, these considerations may be of interest for measurements in environments of different refractive index to water, such as those containing glycerol or sucrose, even though such refractive index tuning is unlikely to have a dramatic impact on the ultimate performance of light-based single molecule detection and mass measurement.

MP requires the removal of a static background image, which is the main reason for generating ratiometric images^1,2,19^. This background resembles a speckle pattern generally attributed to nanoscale roughness of microscope cover glass. Indeed, a recent report successfully correlated nanoscale roughness measured by atomic force microscopy with the corresponding image contrast^20^. Using the reported surface roughness parameters in terms of lateral (∼ 100 nm) and vertical (∼ 2 nm) dimensions, we simulated the resulting phase retardation *ψ* of the reflected field, through:

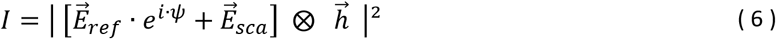

Note that *ψ* describes an effective phase change, which also accounts for the phase delay of the electric field that transmits through the glass-water interface and eventually leads to protein scattering. To obtain the speckle-like appearance in our simulation we create a spatial array of randomly chosen numbers drawn from a uniform distribution 𝒰 _[−1,1]_, which is then spatially low-pass filtered to yield a near diffraction-limited speckle pattern. Overall, this yields a surface height map Δ*h* (varying between ± 0.8 nm^20^), which can be converted into the relevant phase distortion (**Fig. 1e**) via:

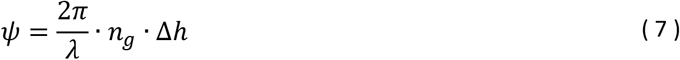

Including this phase variation in our model predicts an image in agreement with experimental results (**Fig. 1f**)^9^, effectively resulting in a locally-varying reflection coefficient.

Given that we can now produce raw images of both microscope cover glass and of single proteins with appropriate optical contrasts and spatial patterns, we can simulate a standard MP experiment (Methods 3 & 4). Here, a cleaned glass coverslip is usually covered by a dilute solution of biomolecules of interest, which bind non-specifically to the glass surface over time. These binding events are best visualised by averaging a series of camera frames of the glass surface and computing the relative difference between consecutive sets as a function of time^3^ (see Methods 5). In this way, individual molecules are revealed, even though their contrast is much smaller than that generated by the glass roughness (**Fig. 2a**).

**Figure 2:**
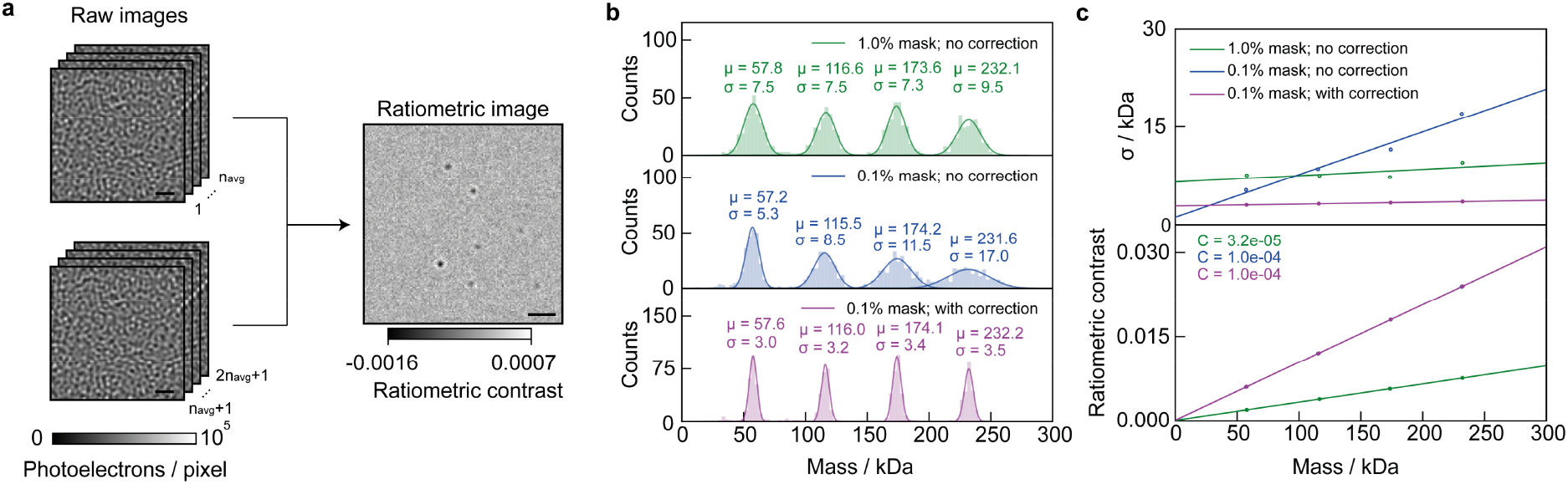
Simulation of protein landing events and resulting mass distributions as a function of mask strength. **a** Two consecutive sets of N frames are averaged before computing the ratiometric image, revealing individual proteins landing at different positions as a function of time (Scale bars: 1 μm). **b** Mass histograms for four oligomeric states of a protein simulated as spheres of radius 3.8 nm and refractive index 1.46 using different mask strengths and local reflectivity correction.^9^ In all cases, the power incident on the detector was kept constant. **c** Standard deviation of the fitted distributions (top) and ratiometric contrast (bottom) as a function of protein mass.

Aside from detection sensitivity, the key performance parameter that determines the utility of MP as an analytical technique is the achievable mass resolution, which originates in the measurement precision achievable on a molecule-by-molecule basis. Using illumination power (1% mask: 0.025 MW cm^-2^; 0.1% mask: 0.25 MW cm^-2^), wavelength (445 nm) and 10^7^detected photoelectrons per pixel (Methods 2), subsequent to temporal and spatial binning, we find peak widths similar to optimal experimental results on the order of *σ* = 8 - 9 kDa (**Fig. 2b**, top). Using a mask with lower transmission in combination with higher illumination power leads to reduced peak widths at low mass, but a broadening as mass increases. This is caused by the influence of the glass roughness on the ratiometric contrast, which now not only depends on the scattering coefficient but also on the locally-varying reflectivity of the glass-water interface (neglecting the purely scattering term):

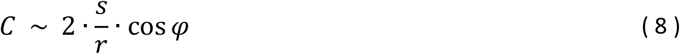

with *cos φ* describing the phase difference between 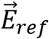 and 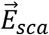 at the detector plane. The glass roughness now results in the ratiometric contrast being dependent on where a particle lands on the glass coverslip. In the simulation, this broadening can be minimised by performing a correction step (based on^21^) by multiplication with 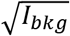, as this removes the dependence on :

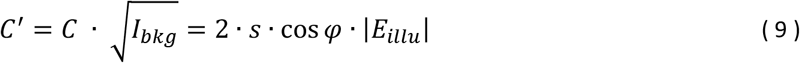

This results in *σ* < 5 kDa peak widths but requires a homogeneous illumination field that is non-trivial to achieve in practice. In all cases, we observe behaviour that agrees with expectations based on the selective reduction of reflected light by the transmission mask. A ten-fold reduction in reflected light is expected to increase the optical contrast by 10^1/2^, which would result in a 3.2-fold reduction in peak width (assuming a constant photon flux reaching the detector), comparable to our results (**Fig. 2c**, top). Similarly, the concomitant increase in the contrast-to-mass conversion factor is also confirmed by our simulations (**Fig. 2c**, bottom). These results validate the potentially high mass accuracy of MP, while assuming a perfectly spherical scatterer.

To explore the effects of molecular shape and orientation beyond the spherical model, we chose double-stranded DNA (dsDNA), which effectively forms linear rods for a few hundred base pairs and below due to the persistence length of DNA^22^. We performed MP measurements of dsDNA (Methods 6) containing 200 and 400 base pairs (bp), while illuminating the sample with circularly polarized light (**Fig. 3a**, top) and observed no significant effect of the elongated shape of the DNA molecule on the ratiometric contrast when compared to proteins of similar mass in terms of peak widths (simulated results agree with our experimental findings and are shown in Supplement 14). The slight broadening for 400 bp DNA likely stems from the fact that the length of the DNA (∼ 136 nm) is no longer negligible compared to the diffraction limit, which can lead to the interferometric signal being spread over a slightly larger PSF and thus lower contrast, ultimately leading to peak broadening. When performing the experiment with linearly polarized light (**Fig. 3a**, bottom), we found a 2-fold increase in peak width, which stems from the underlying variability of contrasts measured on a molecule-by-molecule basis (also see Supplement 14). These results suggest that scatterer shape and orientation can play an important role when employing linearly polarized illumination coupled with fixed molecular orientations. At the same time, it shows that the use of circularly polarized light makes MP essentially insensitive to molecular shape.

**Figure 3:**
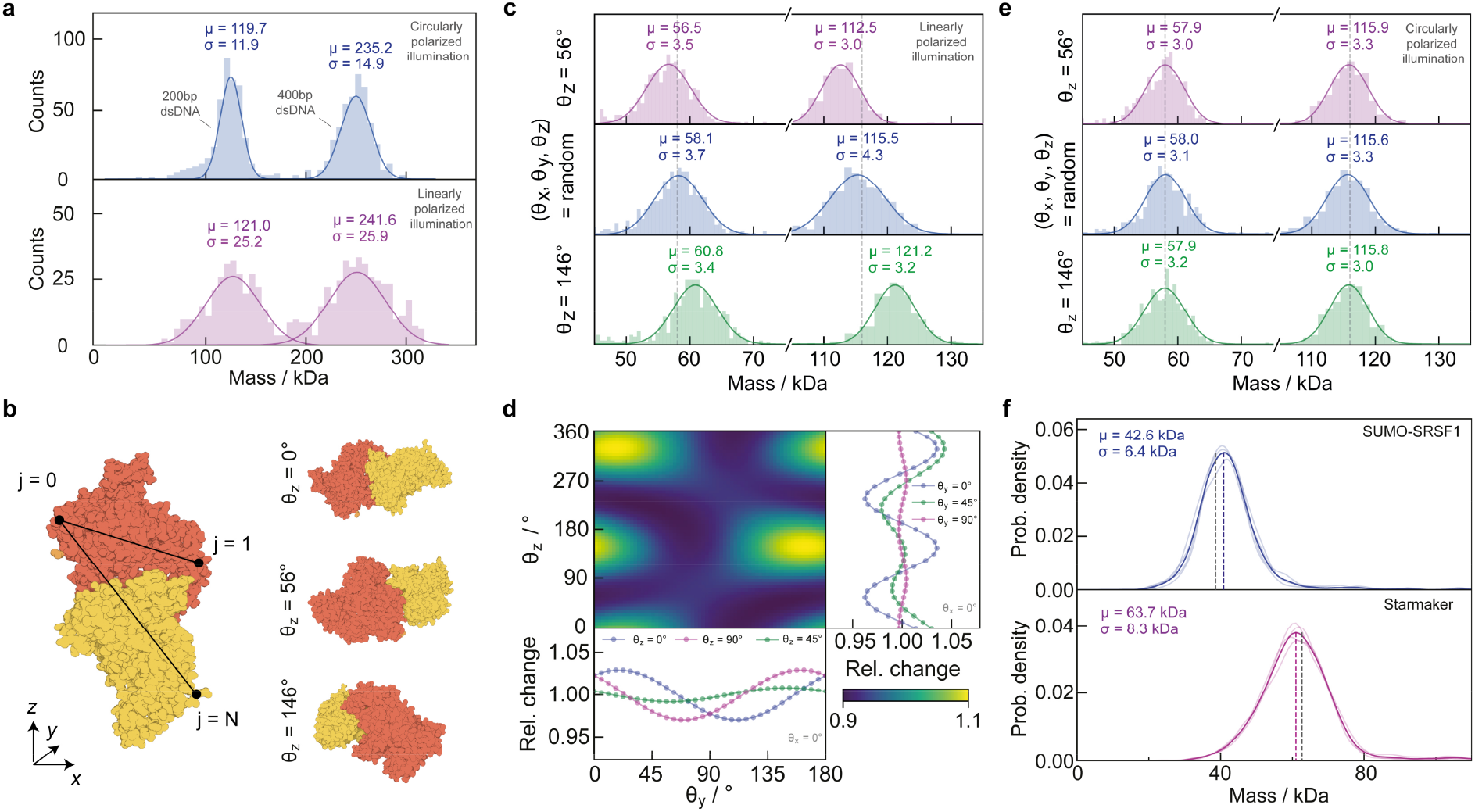
Dependence of mass measurement on biomolecular shape and illumination. **a** Experimentally observed mass distributions for double-stranded DNA illuminated with circularly (top) and linearly (bottom) polarized light. **b** Modelling the shape and orientation of a protein, here the dimer of BSA (PDBID 3V03), by computing the corresponding polarizability tensor. **c** Mass histograms for the BSA monomer & dimer when illuminated with linearly polarized light for two fixed orientations, and for random orientation. **d** Relative change of the ratiometric contrast for the BSA dimer for different orientations relative to the incident polarization. **E** The same simulation as in **c**, except for circularly polarized illumination. **f** Kernel density estimation (Gaussian; bandwidth = 2 kDa) of experimentally measured distributions of partially (SUMO-SRSF1; m = 39.8 kDa; 4 repeats) and fully disordered (Starmaker; m = 66 kDa; 4 repeats) proteins.

While folded proteins do not exhibit the degree of anisotropy as short DNA strands, they are also not spherical, especially in the context of oligomerisation. We therefore turned to a recently reported approach^23^ using atomically-resolved protein structures to compute the polarizability tensors of proteins from pairwise distances of all atoms in the molecule reported in the respective PDB structure (**Fig. 3b**, left; also see Supplement 7 - 8). The resulting polarizability tensor is a 3 × 3 matrix that encodes the anisotropic scattering of a particle, as it connects any specific illumination direction with the scattering in all directions (denoted by *x, y* and *z*):

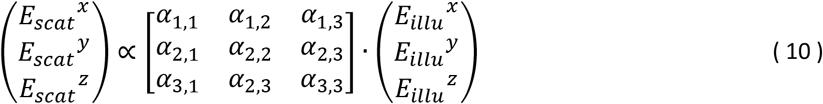

Note that any orientation of the protein can be achieved by multiplying the corresponding rotation matrices ***R***= ***R***_*z*_ **· *R***_*y*_ **· *R***_*x*_ to determine the polarizability tensor ***α***_*rot*_ in the new reference frame^24^:

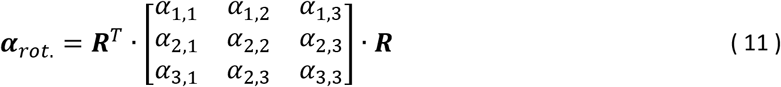

In the first instance, we simulated a series of landing events, where we fixed the orientation of BSA (major axis misaligned: θ_z_ = 56° or aligned: θ_z_ = 56° + 90° = 146°; with respect to the linearly polarized illumination) and compared the resulting mass distributions to those obtained from randomly oriented molecules, for both monomers and dimers (**Fig. 3b**, right). We find significant deviations from the nominal mass in both cases, on the order of 4% of the expected mass for the monomer and dimer (**Fig. 3c**). A deviation of 4% amounts to the maximum observed difference when sampling the full range of possible protein orientations (**Fig. 3d**). Repeating the simulation for BSA with circularly polarized light (**Fig. 3e**) exhibited a drastically reduced dependence on protein orientation upon landing, now amounting to ≪ 1% of protein mass. While the deviations observed for linearly polarized illumination lead to broadening of individual mass peaks, mass photometers reported to date largely rely on circularly polarized light^3^, making these measurements basically insensitive to protein shape. Note that the simulated protein mass for BSA of 58 kDa is lower than the mass based on its amino acid sequence (66 kDa), because the available PDB structure does not contain all atoms. When using the structure of BSA predicted by AlphaFold^25^ (UniProt P02769), we find excellent agreement between the mass predicted by our simulation (64.3 kDa; shown in Supplement 13) and the AlphaFold mass (64.4 kDa).

In addition to protein orientation, the degree to which a protein is folded could also have a substantial effect on the relationship between optical contrast and mass through various factors such as amino acid density or the association of water and counterions, all of which affect the molecular polarizability. We therefore turned to partially and fully unfolded proteins and compared the measured mass using folded proteins as a mass calibrant to the expected mass (Methods 7). SUMO-tagged serine- and arginine-rich splicing factor 1^26^(SUMO-SRSF1) is an 39.8 kDa protein composed of an 11 kDA SUMO-tag, two 8 - 9 kDa structured domains (RRM1/2) and three intrinsically disordered domains totalling 10-11 kDa, making it 25% disordered by mass. The small ubiquitin-related modifier (SUMO) tag is a N-terminal carrier protein that promotes protein folding and stability, allowing for easier production of the desired protein^27^. Using a folded, oligomeric protein as a calibrant (dynamin-1 ΔPRD), we obtain a mass of 42.6 ± 0.64 kDa, in good agreement with the predicted mass (**Fig. 3f**, top) and within the error found for various folded proteins (**Fig. 4b**). We then turned to Starmaker, an 66 kDa, fully disordered protein^28^. Due to its high negative overall charge, Starmaker does not bind adequately to standard microscope cover glass. We therefore functionalised the cover glass (3-aminopropyl)triethoxysilane (APTES) to create a positively charged surface, obtaining a mass of 63.7 ± 0.4 kDa (**Fig. 3f**, bottom), again in excellent agreement with the expected mass. In addition, in both cases we do not see any significant increase in the peak widths compared to folded proteins of similar mass.

**Figure 4:**
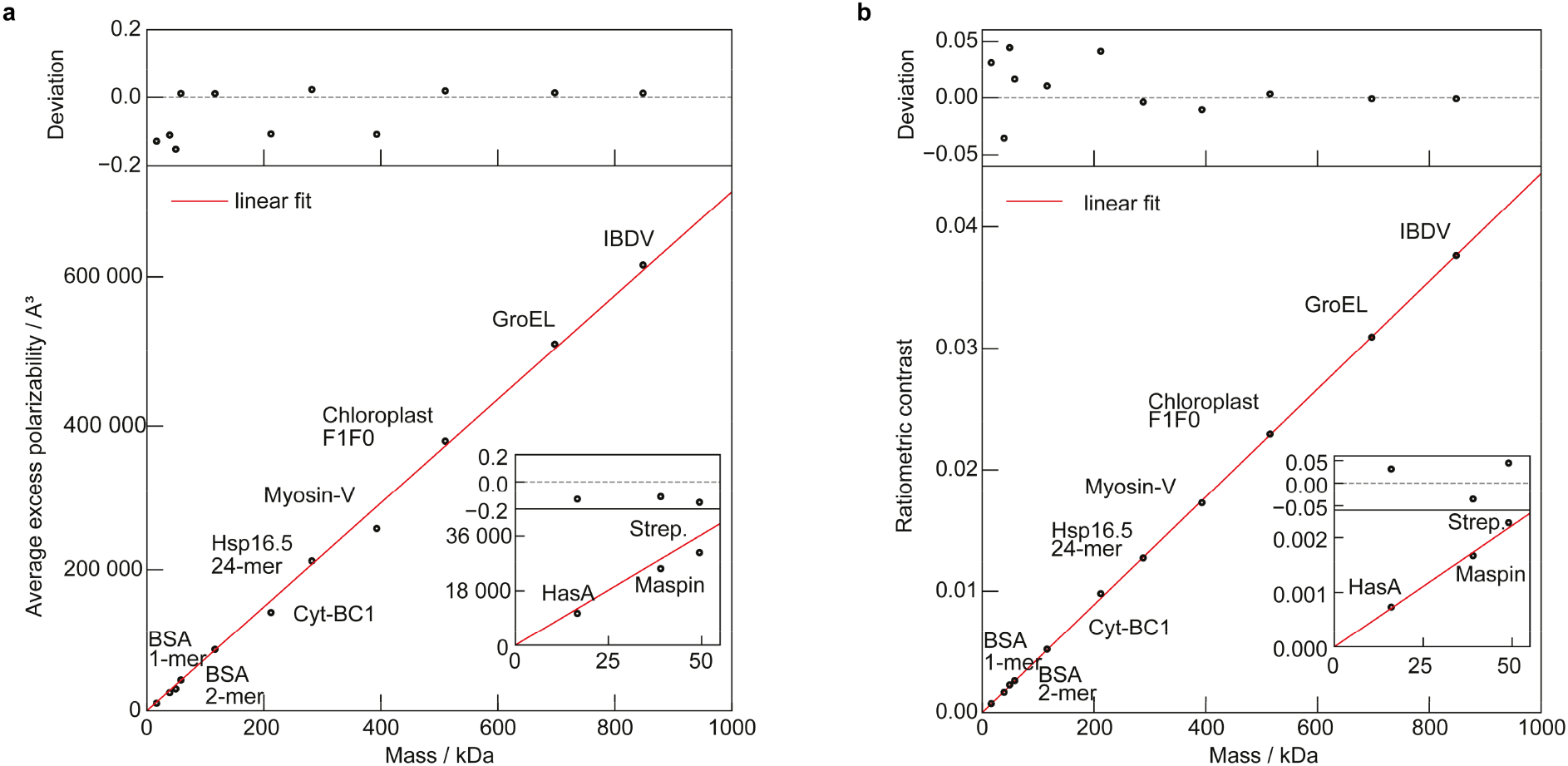
Mass scaling with molecular polarizability and image contrast. **a** Average excess polarizability, and **b**, calculated ratiometric contrast for proteins of mass 10 – 1000 kDa. Slope of linear fit (red line): 724 Å / kDa (a) and 4.4 × 10^−5^ / kDa (b). PDB IDs: HasA = 1B2V; Maspin = 1XQJ; Strep. = 4BX6; BSA = 3V03; Cyt-BC1 = 1BE3; Hsp16.5 = 1SHS; Myosin-V = 2DFS; Choroplast F1F0 = 6FKI; GroEl = 1GR5; IBDV = 2GSY.

We can now explore the previously reported linear relationship between optical contrast and mass for a variety of proteins, using random orientations for landing events and circularly polarised illumination as in the experiment. We find that the resulting (average) excess polarizability^16^ scales linearly with the respective molecular mass, derived from the amino acid sequence in the respective PDB entry (**Fig. 4a**; the respective PDB IDs are given in Methods 8, Table 1), with the polarizability change per mass *δ α*:

**Table 1:**
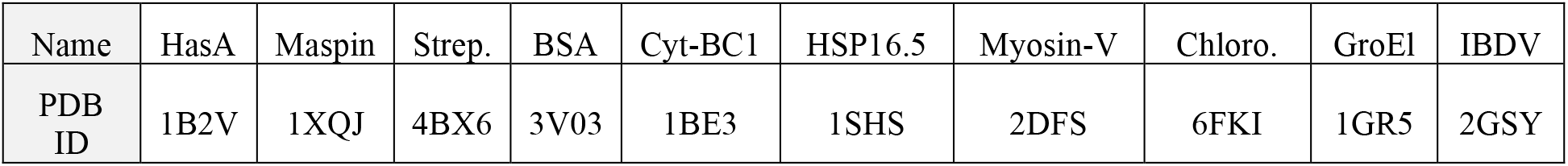
PDBIDs for the different proteins used to investigate the linear relationship between protein mass and scattering strength (see Fig. 4 a & b).

| Name | HasA | Maspin | Strep. | BSA | Cyt-BC1 | HSP16.5 | Myosin-V | Chloro. | GroEl | IBDV |
| --- | --- | --- | --- | --- | --- | --- | --- | --- | --- | --- |
| PDB ID | 1B2V | 1XQJ | 4BX6 | 3V03 | 1BE3 | 1SHS | 2DFS | 6FKI | 1GR5 | 2GSY |

**Table 2:**
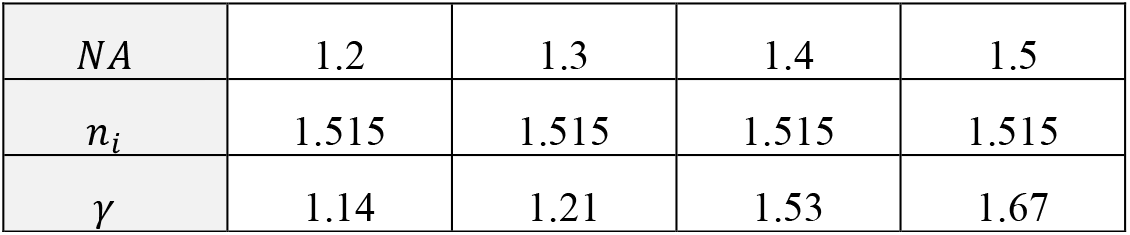
Enhancement factor describing the scattering of a dipole at the glass-water interface for different NA.

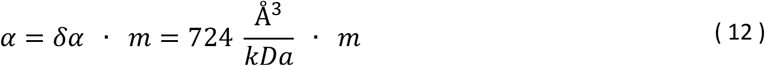

which is slightly larger than that computed from bulk refractive index measurements^4^ (460 *Å*^3^ kDa^-1^). This deviation might be (partially) explained by including the factor *n*_*m*_^2^ into the absolute polarizability value, whose definition dependents on how the scattering process is introduced in the corresponding calculation. When computing *α* for a range of proteins, some exhibit substantial deviation (>15%) from the expected linear relationship, especially below 200 kDa. To determine whether these variations in polarizability are reflected in real MP measurements, we used the full polarizability tensor model to calculate the respective ratiometric contrast from simulated landing assay movies as a function of protein mass. We found a linear relationship, this time with an RMS error of 2.4% and improved agreement above 200 kDa (**Fig. 4b**), both of which agree well with experimental results^3,9^. The most likely reason for this improvement is that the average of the polarizability tensor does not report the scattering response to circularly polarized light, but rather is an average measure of the particle scattering strength^23^. We emphasise that we used the expected mass as calculated from the atomic positions in the respective PDB file rather than the nominal protein mass for all calculations, meaning that most proteins are found below their mass expected from their amino acid sequence.

Equipped with a realistic and quantitative description of the biomolecular polarizability and image formation, we can now deduce a general equation of the achievable signal-to-noise ratio (SNR) as a function of the key experimental parameters. First, we derive the number of detected, *scattered*, photons (*N*_*Sca*_) per pixel and exposure time as (for the full derivation, see Supplement 10):

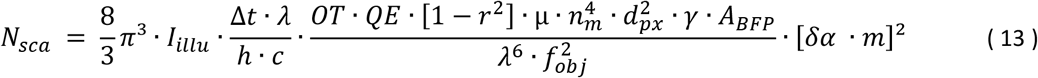

with the following parameters defined in SI base units: *I*_*illu*_: illumination intensity [W/m2]; Δ : effective exposure time per image [s]; *λ*: laser wavelength [m]; *h*: Planck’s constant [Js]; *c*: speed of light [m/s] *OT*: optical throughput; *QE*: quantum efficiency; *r*^2^: reflectivity of glass – water interface; *μ* = *sin*^−1^(*min* [*NA*/*n*_*i*_, 1])/ *π*: collection efficiency of the detection objective; *d*_*px*_ : effective pixel size in sample space [m]; 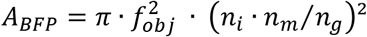: area of the accessible back-focal-plane, limited by the critical angle [m2]; *γ*: enhancement factor due to aplanatic factor & scattering beyond the critical angle (e.g. *γ* ∼ 1.58, for a 1.42 NA oil-immersion objective; see Methods 9 and Supplement 10); *f*_*obj*_: focal length of detection objective [m]; *n*_*m*_: refractive index of buffer medium; *m*: mass of protein [kDa]; *δ α*: polarizability per kDa, i.e. the slope in Fig. 4a.

The detected, *reflected*, number of photons *N*_*ref*_ (per pixel and exposure) is similarly given as:

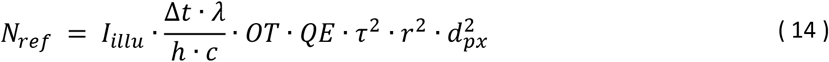

with *τ*^2^ indicating the (power) transmission coefficient of the mask in the BFP. The shot-noise limited *SNR* in terms of the ratiometric contrast follows then as (see the Supplement 10):

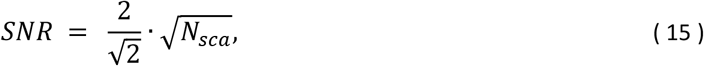

with the factor of two originating from the interferometric nature of the signal and the 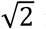 from the comparison of two subsequent set of frames, required to remove the static background and form the ratiometric image. This can be converted into an equivalent expression for the detection limit of MP, i.e., the smallest detectable mass *m*_*q*_ at a certain *SNR* level. For this we set *SNR* = *q* and solve for *m*:

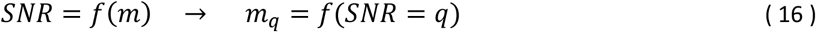

With *f* denoting a functional dependency. Similarly, we can define a measure describing the lowest achievable mass resolution *σ*_*m*_, the (Quantum-) Cramer-Rao-Lower-Bound as defined in ^29^:

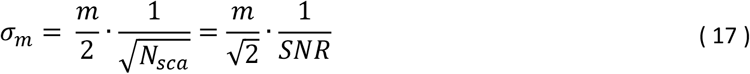

We also find that this optimum achievable mass resolution is directly linked to the SNR-equivalent mass, with 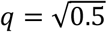 (see Supplement 10 for the derivation):

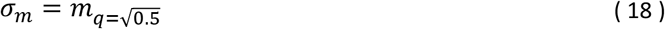

Meaning that the highest attainable mass resolution is equivalent to the mass that achieves an 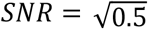 in the shot-noise limited regime.

Given that most of the parameters are essentially fixed in realistic experimental scenarios, or only vary marginally as a result of the details of experimental implementation, we can simplify these expressions to depend only on some key experimental details, specifically illumination power, exposure time, wavelength and molecular mass (in kDa).

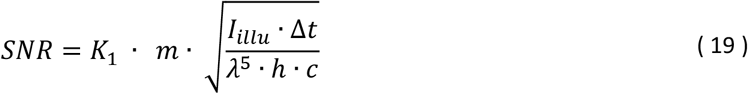

with 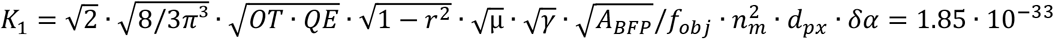 in units of *m*^*4*^/*kDa*, assuming: *OT*= 0.8; *QE* = 0.^7^0; *r*^2^ = 0.00*4*; *f*_*obj*_ = 3 mm; *δ α* = ^7^2*4 Å*^3^/*kDa*; *n*_*m*_ = 1.333; *μ* = 0.*4*0; *γ* = 1.58; *A*_*BFP*_ = 50 mm2; *d*_*px*_ = 80 nm. Yielding an *SNR* ∼ 21; *m*_*q*=3_ ∼ 9.5 kDa and *σ* _*m*_ ∼ 2.2 kDa for *I*_*illu*_ = 0.1 MW cm^-2^, Δ*t* = 100 ms, *λ* = *44*5 nm and *m* = 66 kDa.

Experimental images, however, including those consisting of buffer medium only, or even ultrapure water, reveal a dynamic, speckle-like background at high imaging sensitivity that cannot be removed by temporal averaging and cannot be attributed to sample drift, here shown by comparing 120 and 480 ms averaging time, plateauing at a contrast on the order of a 5 kDa protein (**Fig. 5a**). This background is likely the current limiting factor to both improving mass resolution, and the absolute detection limit of mass photometry, and is close to that reported recently using machine learning^30^. We currently have no clear explanation as to the origin of this additional noise source. As a first attempt we include it into our (SNR-) model, by adding it in quadrature to the shot-noise variance and find for the SNR including additional excess noise *SNR*_*exc*._ (with *σ*_*exc*._ being the added variance in terms of photon counts):

**Figure 5:**
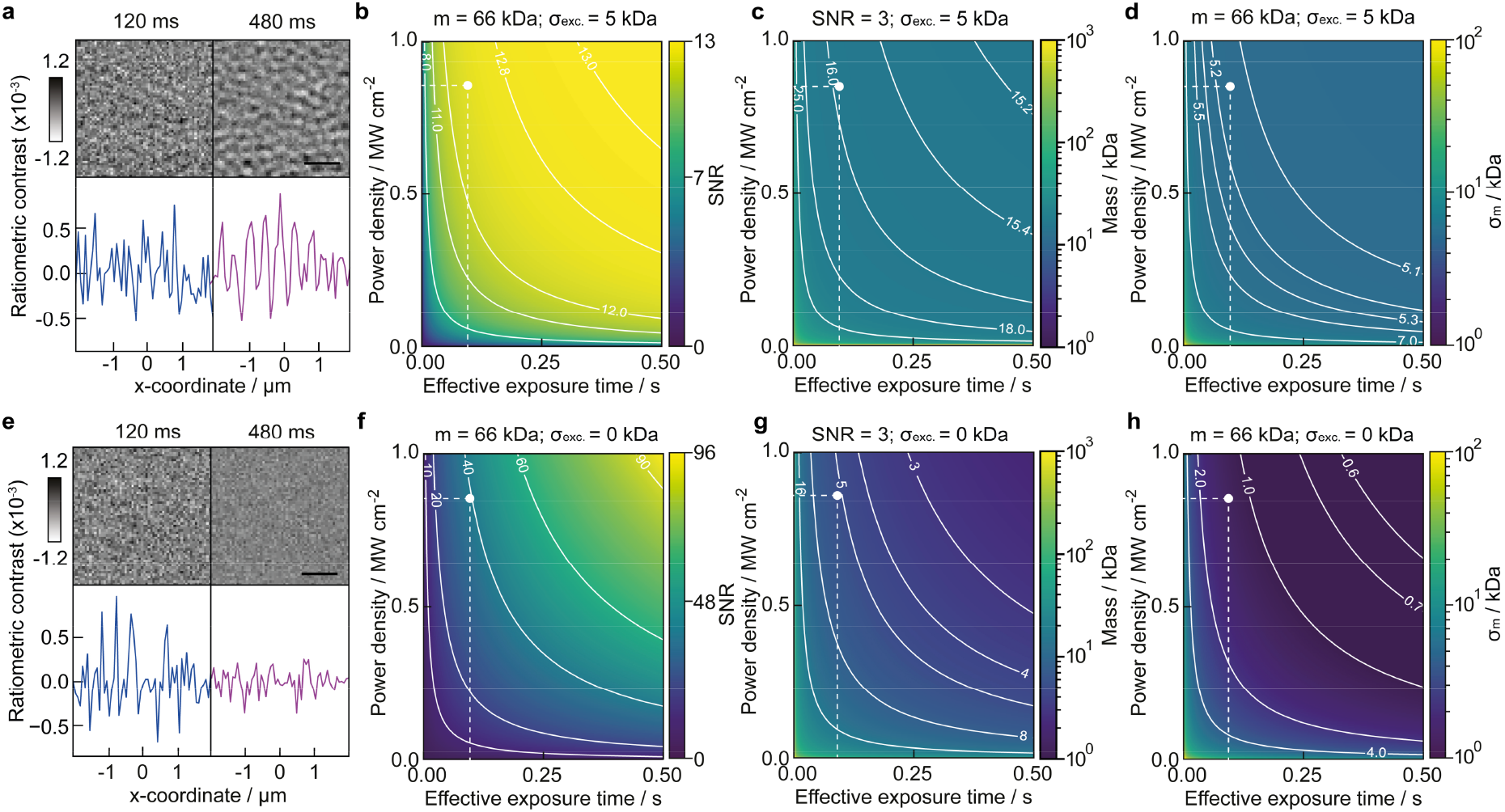
Current and future limits of optical mass measurement of single biomolecules. **a** Ratiometric images of buffer medium-only (scalebar = 1 μm) for different integration times. **b** Achievable signal-to-noise ratio when detecting the BSA monomer (m = 66 kDa). **c** Smallest detectable mass m_q=3_ at SNR = 3. **d** Mass resolution a_m_. All given as a function of effective exposure time and illuminating power density in the presence of excess noise on the order of 5 kDa. **e - h**, Same images and dependencies for purely shot noise limited performance. The white dots indicate experimental parameters from ^3^.

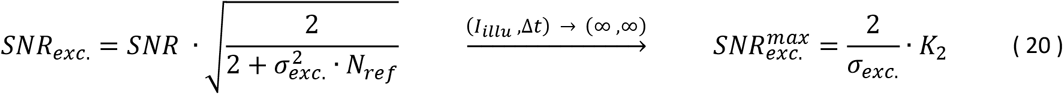

and the constant 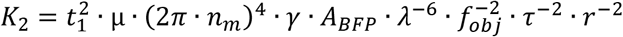. Note that *SNR*_*exc*._ Naturally results in shot-noise limited performance for 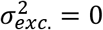 and yields a finite, maximum, 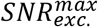. The added influence of *a*_*exc*._ has an overall impact on key performance parameters for protein detection and characterisation, such as the achievable SNR (**Fig. 5b**; with 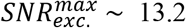), the lowest mass detectable (**Fig. 5c**), and the achievable mass resolution (**Fig. 5d**).

We can now compute both theoretically achievable performance in terms of SNR, mass resolution and detection limit (assumed for SNR = 3) both in the presence (**Fig. 5a - d**) and absence (**Fig. 5e - h**) of a non-shot noise contribution amounting to 5 kDa RMS as currently observed experimentally. For realistic simulations, we find good agreement with previous reports^3^, such as an SNR of ∼ 12.5 for BSA at an exposure time of 100 ms and 0.85 MW cm^-2^. Similarly, the recently reported SNR of 1.4 for a 9 kDa protein^30^ appear realistic, given that we find *m*_*q*=1_ ∼ 5.2 kDa with and ∼ 1.7 kDa without the additional baseline noise (for the same exposure time and illumination power density; see Supplement 16). Overall, our simulations in the absence of excess noise demonstrate that significant improvements in performance are still achievable with realistic illumination powers (∼ 1 MW cm^-2^) and exposure times (< s), such as 1 kDa mass resolution and few kDa detection sensitivity, even in the absence of advanced image processing.

## Discussion

We have presented a numerical approach that enables us to compute the optical contrast generated by individual biomolecules based on their atomic structure, orientation and shape. Our results compare well with experimental data obtained by MP, suggesting that our model is indeed quantitative. We find a clear dependence of optical contrast on molecular shape in extreme cases such as short DNA strands illuminated by linearly polarized light, that weakens for folded proteins, and is essentially eliminated when using circular polarization. The predicted mass accuracy on the order of 2.4% RMS matches that observed experimentally (2%), as does the computed dependence of optical contrast on the strength of the attenuation mask used. Our results on intrinsically disordered proteins support the hypothesis that the major determinants for the molecular polarizability are the constituent amino acids. In terms of achievable sensitivity, we present evidence for a dynamic, speckle-like background with a signal magnitude comparable to a 5 kDa protein. This background currently limits the ultimately achievable detection sensitivity, and also affects the achievable mass resolution. The resolution is further affected by the static speckle-like background caused by microscope cover glass roughness, although it can be corrected computationally.

Our results provide a quantitative framework for rationalising label-free optical detection of single biomolecules. Polarizabilities, realistic incident power densities and detection efficiencies effectively define achievable detection sensitivity and measurement precision at the single molecule level, which translates into mass resolution. We emphasise that these relationships are independent of the optical approach, whether using total internal reflection^5^ or scattering from nanochannels^4^. While there will be subtle differences in the achievable power densities and detection efficiencies, the presented limits are likely to be highly representative of what can be achieved in terms of mass measurement using light-based detection of single biomolecules. In all cases, when comparing experimental with these theoretical results, it is essential that any non-shot noise contributions to image background are considered and quantified carefully, given their influence on measurement sensitivity and precision.

What is most encouraging, however, is that there appears substantial scope for improvement that will enable quantitative characterisation of complex mixtures of biomolecules with a resolution and sensitivity that covers almost all species and interactions. Moreover, implementation of approaches that enable extended observation of individual molecules either in nanochannels^4^ or on bilayers^13,31^ could bring about even further improvements, much in the spirit of the advances brought about by similar strategies in mass spectrometry, such as charge detection and orbitrap mass spectrometry^32,33^. Alternatively, if high resolution and sensitivity is not required, integration times can be drastically reduced, which will enable measurement at higher analyte concentrations, providing access to a broader range of affinities and ultimately weak interactions, further broadening the application scope of MP for characterising biomolecular interactions and dynamics.

## Methods

### 1 Description of general image formation

We describe the image formation in MP as a 3D (complex-valued) convolution of electric fields (reference and scattered) with an amplitude point-spread-function (APSF)^24^, which is implemented as a multiplication in Fourier space (through the *convolution* theorem). The strength of the reference and scattered fields are given by Fresnel’s coefficients *r* & *t* (reflection and transmission) at a glass-water interface and by the scattering coefficient of a small spherical particle, in the Rayleigh regime^14^. The scattering coefficient *s*, is related to the polarizability *α*, i.e., to the volume *V* of the spherical scatterer and the refractive indices of the protein *n*_*p*_ and the surrounding medium *n*_*m*_. High numerical aperture focusing effects are modelled using a geometric transformation^34^, which also includes the aplanatic factor^24^ and the additional Gouy phase shift of the scattered light^35^ (all included in the 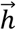 -term). The influence of phase aberrations due to the refractive index mismatch at the coverslip interface is described using the Gibson-Lanni model^36^. The scattering of the protein at the water-glass interface is given as a distribution at the back-focal plane of the objective^37^, which further enables us to add a mask that attenuates and/or delays the reference component^9^. A more detailed description of the underlying theoretical framework is given in Supplements 1 - 8.

### 2 Simulation parameters

All numerical results assume a 1.42 *NA* oil-immersion (*n*_*i*_ = 1.515) objective while imaging at a wavelength of *λ* = 445 nm. The data shown in Fig. 1 corresponds to an effective pixel size of 0.057 μm with 70 × 70 pixels, yielding a field-of-view (FoV) of 4 μm × 4 μm = 16 μm2. The step size along the axial direction was chosen to match the lateral size, ± 2 μm propagation from the nominal focus. In case of the simulated landing assay data (all remaining figures) the effective pixel size was 70 nm at 128 × 128 pixels, resulting in a FoV of ∼ 80 μm2, while only computing a single in-focus slice. In this case we set the attenuation mask to add a π/2 phase delay for 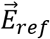, which results in phase-matching of the two fields at the detector, resulting in optimum contrast at the nominal focal plane (see Supplement 5 & 12). The illumination was set to be (right-) circularly polarized, except for Fig. 3b,c were we defined it to be linearly polarized. In terms of illumination intensity 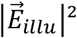, we directly defined the detected photons at the detector, omitting the need to specify the overall efficiency of the detection system (including quantum efficiency of the detector, losses at the optics, …). For the landing assay simulations this yielded in 10^6^ photons per 70 nm pixel per 500 μs exposure time, which corresponds to the 10^7^ photoelectrons mentioned earlier, when performing the ratiometric calculation with *N* = 100.

### 3 General measurement routine

Data was either collected on a TwoMP (Refeyn, UK) or on a custom-built mass photometer. Coverslips (Menzel-Glaeser, 24 × 50 mm # 1.5 SPEZIAL; Thermo Fisher Scientific, U.S.) were cleaned to remove any contaminants by sonication for 5 minutes in 50/50 isopropanol and Milli-Q water (Merck, Germany), followed by 5 minutes in Milli-Q only. They were then dried using N_2_ and stored in a covered box to prevent re-contamination. Immediately before measurement, a silicon gasket (Grace Bio-Labs CultureWell, 3 × 1 mm; U.S.) was laid on the coverslip to contain the sample. The coverslip was placed on a sample-stage (xyz for TwoMP; z-only for custom-built system) above the objective and a small amount of immersion oil (Zeiss, Immersol 518 F; Germany) was added between the coverslip and objective to form a continuous interface. Once the gasket and objective were aligned, buffer was loaded into the gasket using a micropipette (Gilson Pipetman, U.S.). In case of the TwoMP, this allowed the focus position of the setup to be found and the autofocus to be set before the protein began binding to the coverslip. The custom-built system was operated without an autofocus but proved to be stable enough over the recording time. Once focus had been set, the protein was diluted in an Eppendorf tube (Eppendorf, 1.5 mL; Germany) to give 20 μL of sample, at a concentration of 10-50 nM. The sample was added to the gasket and the focus quickly rechecked. If the added sample had been kept on ice, the refractive index of the solution could change when the preloaded buffer and sample were mixed due to the difference in temperature, which changed the focus position slightly. A movie was then recorded. The protein dynamin-1 ΔPRD was used as a mass calibrant. It is highly stable, easily produced in large volumes and oligomeric, providing a large number of species of known mass (sometimes up to 7) with which to calibrate, increasing the accuracy of the calibration.

### 4 Materials

Reagents used were from Sigma-Aldrich (U.S.), unless otherwise stated. Water was ultrapure Milli-Q (Merck, Germany) and all solutions were filtered through a 0.2 μm filter (Millipore, U.S.) before use.

The disordered protein SUMO-SRSF1 containing the solubilising mutations Y37S and Y72S^38^ was cloned by Gibson assembly into a pET28 plasmid (kind gift of B. B. Kragelund, University of Copenhagen, Denmark) downstream of a His_6_-SUMO tag. The construct was transformed into chemically competent C41 *E. coli* (Lucigen). Cultures were grown in 2xYT medium supplemented with 50 μg/ml kanamycin at 37°C until an OD_600_ of ∼1.5 was reached, and protein expression was induced using 0.5 mM IPTG at 20°C overnight. Cells were harvested, resuspended in buffer A (20 mM sodium phosphate pH 7.5, 800 mM NaCl, 5% glycerol, 0.01% Tween-20, 2 mM dithiothreitol (DTT), 150 mM L-arginine, 150 mM L-glutamate) supplemented with 10 mM MgCl_2_, 10 U/ml benzonase (Merck) and 20 μg/ml RNase A (NEB), and lysed by passing the suspension three times through a high-pressure homogeniser (HPL6, Maximator) cooled to 4°C at 15,000-20,000 psi. The lysate was clarified by centrifugation and applied to a HisTrap Excel column (Cytiva, 5 ml per 1 l cell culture) equilibrated in buffer A. The column was washed with 15 column volumes (CVs) of buffer A, followed by 7 CVs buffer B (20 mM sodium phosphate pH 7.5, 3 M NaCl, 5% glycerol, 0.01% Tween-20, 2 mM DTT, 100 mM L-arginine, 100 mM L-glutamate) and 7 CVs of 97% buffer A and 3% buffer C (20 mM sodium phosphate pH 7.5, 800 mM NaCl, 5% glycerol, 0.01% Tween-20, 2 mM DTT, 150 mM L-arginine, 150 mM L-glutamate, 500 mM imidazole), before elution with buffer C. Fractions containing protein were pooled and diluted at least 8-fold with buffer D (20 mM HEPES pH 8.0, 10% glycerol, 0.001% Tween-20, 0.5 mM Tris (2-carboxyethyl) phosphine, TCEP) and 5 M NaCl until the sample was clear (∼0.8 M ionic strength or 55 mS/cm). Nucleotides and protein contaminants were removed from the sample by ion exchange chromatography using a MonoS 5/50 GL column (Cytiva) and a gradient of 30-70% of buffer E (20 mM HEPES pH 8.0, 2 M NaCl, 10% glycerol, 0.01% Tween-20, 0.5 mM TCEP). Fractions with absorbance ratios of A260/A280<0.7 were pooled, flash frozen in liquid nitrogen and stored at -80°C. Starmaker was prepared as described previously^28^.

Stock solutions of SUMO-SRSF1 were at 29 μM protein. 1 μM aliquots were prepared in 20 mM HEPES (pH 7.4), 1 M NaCl, 1 mM DTT, 5% glycerol buffer and flash frozen. High salt was required to stabilize the protein. DTT is a reducing agent, preventing oligomerisation via disulfide bonds as SUMO-SRSF1 contains two internal cysteines. Starmaker was diluted in 20 mM Tris (pH 8.4), 50 mM KCl. The concentration was unknown, so measurements of a range of dilutions from the original stock were taken to estimate the concentration. For Starmaker, coverslips were positively charged with APTES. Coverslips were cleaned as described before, then plasma cleaned for 8 minutes. The coverslips were washed in acetone and submerged in a 2% APTES/acetone solution. After 2 minutes, the coverslips were washed with acetone again and placed in an oven for 1.5 hours at 110 °C. Finally, the coverslips were sonicated for 5 minutes in isopropanol, then water and dried under N2.

### 5 Data analysis

For the analysis of the recorded data, we used an in-house python package. Raw frames from the measurement *I*_*i*_ were imported and converted into ratiometric frames, by averaging two stacks of *n*_*avg*_ raw frames; *Ī*_1_ from frame (*i*) to (*i* + *n*_*avg*_) and *Ī*_2_ from frame (*i* + *n*_*avg*_ + 1) to (*i* + 2*n*_*avg*_ + 1):

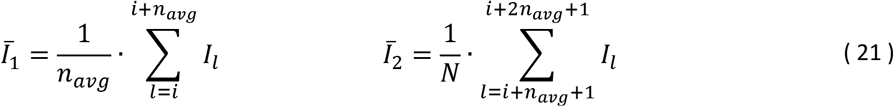

Those two stacks are then used to compute the relative difference:

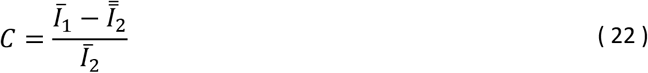

This eliminates the background signal (from the glass roughness) that is constant throughout the measurement. Protein binding events that occur during the measurement are not constant, hence are not removed when computing the relative difference. They appear as spots that fade in and out of the image as the protein binds and then becomes part of the background. Once the ratiometric frames have been calculated, protein binding events are identified by filtering the detected events for groups of pixels that meet a minimum radial symmetry and for pixels with a minimum signal above the background noise.

A PSF (either a theoretical^3^ or experimental model) is then fitted to each event to determine its contrast. The use of an experimental PSF is necessary when analyzing data that corresponds to illuminating the scatterer with linearly polarized light (e.g. in the DNA measurement). To obtain the experimental PSF, the initial PSF detection parameters (found with the theoretical model) are used to align the cropped images (typically 7 × ^7^ pixels) of the found landing events, by employing a cubic spline interpolation. Those sub-images are then averaged together, whilst outliers (Pearson’s correlation test) are being removed from this average. Then the resulting cubic spline model is used to determine the contrast of each landing event. The obtained contrast values for all events are then plotted as a mass histogram. For a particular species, the event contrasts are usually normally distributed allowing a Gaussian to be fitted to the respective peaks. The Gaussian position, width and area were used to characterize the contrast of each peak *μ*, standard deviation *σ* and counts, respectively. To generate a mass calibration, this procedure was applied to a measurement of dynamin-1 ΔPRD. The peak contrasts were used for calibration by plotting against the corresponding species mass. In case of the simulated data, the calibration was performed against the first four oligomeric states of BSA shown in Fig. 2 b, while deliberately reducing the numerically applied shot noise to obtain an accurate calibration.

### 6 Measurements of the double stranded DNA

Data was taken on both a custom-built MP setup that uses 465 nm linearly polarized illumination and a TwoMP (Refeyn, UK) with 488 nm circularly polarized illumination. APTES coverslips with silicone gaskets were prepared via the procedure described above. 200 and 400 base pair double stranded DNA were prepared using standard procedures^39^. For the MP measurements, 200 bp dsDNA and 400 bp dsDNA were diluted to 3 nM and 4 nM respectively in Dulbecco’s phosphate buffered saline. 20 μL of the sample was added to a gasket containing 5 μL of buffer. Data was acquired for 120 s following sample refocusing. The contrast values were converted into mass using a calibration curve (obtained from a measurement of dynamin-1 ΔPRD).

For the custom-built linearly polarized MP setup, data was acquired with the following parameters: 959 μs exposure time, 787 fps, 3.5 × 11.7 μm field of view, 3 × 3 pixel binning and 4-fold temporal averaging. For the TwoMP, data was acquired with the following parameters: 1380 μs exposure time, 699 fps, 2.7 × 10.9 μm field of view, 6×6 pixel binning and four-fold temporal averaging. Both datasets were analyzed such that the ratiometric window size amounted to ∼ 50 ms.

### 7 Measurements of partially and fully unfolded proteins

Data was taken using a TwoMP mass photometer (Refeyn, UK) and analyzed using a custom-written Python package, based on the procedure described in Young et al. ^3^. Coverslips (Menzel-Gläser, 24 × 50 mm # 1.5 SPEZIAL; Thermo Fisher Scientific, U.S.) were cleaned, a silicon gasket (Grace Bio-Labs CultureWell, 3 × 1 mm; U.S.) was laid on top and 4 μL of buffer medium was added,. Next, the protein was diluted in an Eppendorf tube (Eppendorf, 1.5 mL; Germany) to give 20 μL of sample and added into the gasket. Movies containing ∼ 1000 – 5000 binding events were recorded (60 s), analyzed and converted into mass using a calibration curve (generated from a measurement of the oligomeric peaks of dynamin-1 ΔPRD).

SUMO-SRSF1 was diluted to 20 nM in a buffer of 20 mM HEPES (pH 7.4), 1 M NaCl, 1 mM DT and 20 mM NaCl. A 20 mM Tris (pH 7.4), 50 mM NaCl buffer was used to dilute Starmaker, allowing for a reduction or an increase in salt concentration upon measurement. For both proteins 4 repeats were taken at each condition, with no significant unbinding in any repeat.

### 8 PDBIDs of several analyzed proteins

A list containing the PDBIDs of the respective proteins shown in Fig. 4 a & b are:

## 9 Enhancement factor describing the scattering near a glass-water interface

Here we report the enhancement factor *γ* that describes the additionally detected scattering which corresponds to the near-field of a scatterer at the glass-water interface. Details on the computation of *γ* are given in Supplement 10.

## Data availability

Data that support the findings of this study are available from the authors upon request.

## Acknowledgments

We would like to thank Dan Loewenthal for creating the 3D rendering of the BSA dimer shown in Fig. 3b, Roi Asor for fruitful discussions regarding obtaining the polarizability tensor from PDB-structures, Magdalena Wojtas and Andrzej Ożyhar for the kind gift of the Starmaker sample, and Antoine Cléry and Andrea Holla for helpful discussions regarding protein preparation and selection, respectively.

This work was funded by European Research Council (ERC) Consolidator Grant PHOTOMASS 819593, the Engineering and Physical Research Council (EPSRC) Leadership Fellowship EP/T03419X/1 (P.K.), a FEBS Long-term Fellowship (M.S.), and the Swiss National Science Foundation (B.S.).

## Author contributions

Conceptualization: J.B. and P.K.; Methodology: J.B., J.S.P., I.C. and P.K.; Investigation – simulation & theory: J.B.; DNA sample preparation: S.H.; Protein sample preparation: M.S.; Investigation - experimental: J.S.P. and I.C.; Data analysis: J.B.; Visualization: J.B. and P.K.; Funding acquisition: P.K., M.S. and B.S.; Project administration: J.B. and P.K.; Supervision: P.K. and B.S.; Writing – original draft: J.B. and P.K.; Writing – review and editing: J.B., J.S.P., I.C., S.H., M.S., B.S. and P.K.

## Supplementary information

Supplementary information accompanies the manuscript.

---

For Table of Contents only

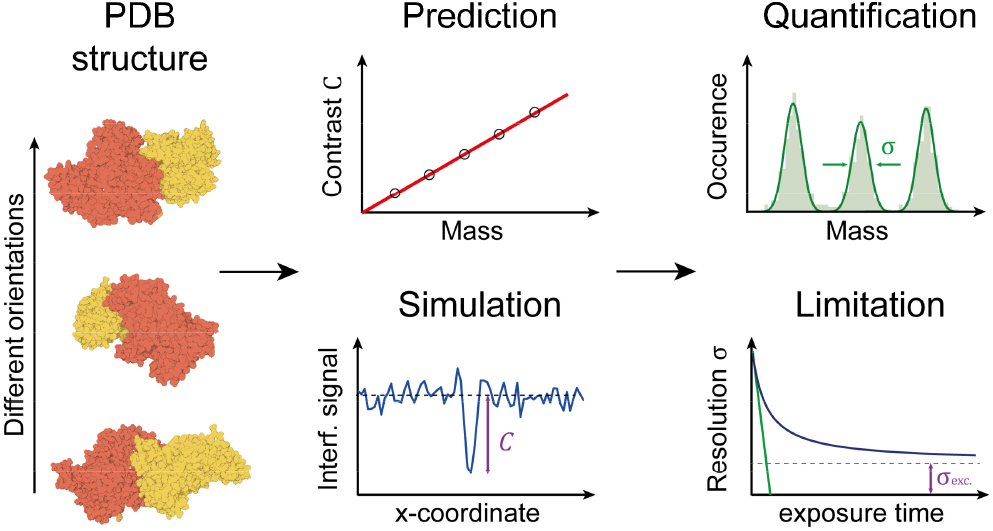

