## Supplemental information for "A quantitative description for optical mass measurement of single biomolecules"


### 1 General image formation model

We model the influence of the optical system (a *widefield* microscope) as a 3D convolution $\otimes$ with an amplitude response function $\mathbf{h}$. The detectable intensity $I$, is then given as [1, 2]:

$$I(\mathbf{r}) = \left|\mathbf{E}_{ref} \otimes \mathbf{h}_{ref}(\mathbf{r}) + \mathbf{E}_{sca}(\mathbf{r}) \otimes \mathbf{h}_{sca}(\mathbf{r})\right|^2 \quad (1)$$

with $\mathbf{E}_{ref}$ and $\mathbf{E}_{sca}$ being the reflected & scattered field at the nominal focal plane (= glass-water interface) and $\mathbf{r} = (x, y, z)^\top$ a vector representing spatial coordinates ($\mathbf{E}_{ref}$ is not spatially depending due to widefield illumination).
Note that the absolute square operation $|\mathbf{E}|^2$ is defined as:

$$|\mathbf{E}(\mathbf{r})|^2 = |E^{(1)}(\mathbf{r})|^2 + |E^{(2)}(\mathbf{r})|^2 + |E^{(3)}(\mathbf{r})|^2 \quad (2)$$

with the superscript $^{(1),(2),(3)}$ denoting the electric field in the $x$, $y$ and $z$ direction. The convolution $\otimes$ is separately performed on each component of the vector fields, e.g. for the first component it is written as:

$$E^{(1)}(\mathbf{r}) \otimes h^{(1)}(\mathbf{r}) = \int_{-\infty}^{+\infty} d\mathbf{v} \;\; E^{(1)}(\mathbf{v}) \cdot h^{(1)}(\mathbf{r} - \mathbf{v}) \quad (3)$$

Which is equivalent to a multiplication in *Fourier* space (convolution theorem):

$$\mathcal{F}\left\{E^{(1)}(\mathbf{r}) \otimes h^{(1)}(\mathbf{r})\right\} = \tilde{E}^{(1)}(\mathbf{k}) \cdot \tilde{h}^{(1)}(\mathbf{k}) \quad (4)$$

with $\mathbf{k} = (k_x, k_y, k_z)^\top$ denoting spatial frequencies and $\mathcal{F}$ being the Fourier transform, defined as:

$$\tilde{E}^{(1)}(\mathbf{k}) = \frac{1}{(2\pi)^{\dim\{d\mathbf{r}\}/2}} \int_{-\infty}^{+\infty} d\mathbf{r} \;\; E^{(1)}(\mathbf{r}) \cdot e^{i\mathbf{k}\cdot\mathbf{r}} \quad (5)$$

with $\dim\{d\mathbf{r}\}$ indicating the dimensionality of the Fourier transform (e.g. $d\mathbf{r} = (dx, dy)^\top \rightarrow \dim\{d\mathbf{r}\} = 2$).

Such a Fourier relationship connects the light distribution at the nominal focal plane with its counterpart in the back-focal-plane (*BFP*) of the microscope objective. When denoting the coordinates in the *BFP* with $\mathbf{r}' = (x', y')^\top$, we see that there exists a geometric transformation $\hat{\mathbf{T}}$ that accounts for the tilting of each light ray towards the nominal focus [3]:

$$\tilde{\mathbf{E}}(k_x, k_y, z = 0) = \hat{\mathbf{T}}(\phi, \theta) \cdot \tilde{\mathbf{E}}(\mathbf{k}') \quad (6)$$

The transformation matrix $\hat{\mathbf{T}}$ is depending on the azimuthal $\phi$ and polar $\theta$ coordinates in the *BFP*:

$$\hat{\mathbf{T}}(\phi, \theta) = \begin{bmatrix} \cos^2\phi \cdot \cos\theta + \sin^2\phi & \sin\phi \cdot \cos\phi \cdot (\cos\theta - 1) \\ \sin\phi \cdot \cos\phi \cdot (\cos\theta - 1) & \sin^2\phi \cdot \cos\theta + \cos^2\theta \\ \cos\phi \cdot \sin\theta & \sin\phi \cdot \sin\theta \end{bmatrix} \quad (7)$$

The real space representation $\mathbf{E}$, at $z = 0$, is then given as an inverse Fourier transform $\mathcal{F}^{-1}$ of each vector component [2]:

$$\mathbf{E}(x, y, z = 0) = \mathcal{F}^{-1}\left\{\tilde{\mathbf{E}}(k_x, k_y, z = 0)\right\} \quad (8)$$

where the (inverse) Fourier transform is applied element wise and $\mathcal{F}^{-1}$ is the complex-conjugate operator of $\mathcal{F}$.

Next, we model the propagation along the optical axis $z$ using the Gibson-Lanni model [4]. It describes the defocusing of an optical imaging system consisting of three layers (immersion, glass coverslip and sample, i.e. buffer). For this we define the optical path difference *OPD*, comparing the usage of the objective in the design case (immersion medium thickness $d_{i,*}$ and refractive index $n_{i,*}$) with the actual experimental scenario (denoted by $d_i$ and $n_i$):

$$OPD(z,z_p) = \frac{n_m \cdot z_p}{\cos\theta_m} + \frac{n_i \cdot d_i(z,z_p)}{\cos\theta_i} - \left[\frac{n_{i,*} \cdot d_{i,*}}{\cos\theta_{i,*}} + n_i \cdot \sin\theta \cdot \left(z_p \cdot \tan\theta_m + d_i(z,z_p) \cdot \tan\theta_i - d_{i,*} \cdot \tan\theta_{i,*}\right)\right] \quad \textbf{(9)}$$

with $z_p$ the axial position of the protein, $n_m$ the refractive index of the buffer medium (here water) and the polar angles $\theta$ in each layer which can be found using *Snell's* law, e.g. $\sin\theta_i \cdot n_i = \sin\theta_g \cdot n_g$. The $z$-dependence is introduced in terms of a variable immersion medium thickness $d_i$ [4]:

$$d_i(z,z_p) = z - z_p + n_i \cdot \left(\frac{d_{i,*}}{n_{i,*}} - \frac{z_p}{n_m}\right) \quad \textbf{(10)}$$

Finally, we obtain the full (3D), real space, electric field distribution by multiplying the *OPD* as a phase factor to the BFP-representation of the respective E-field:

$$\mathbf{E}(\mathbf{r}) = \mathcal{F}^{-1}\left\{\mathcal{F}\{\mathbf{E}(x,y,z=0)\} \cdot G(z,z_p)\right\} \quad \textbf{(11)}$$

with the phase factor $G$ given as:

$$G(z,z_p) = e^{i \cdot \frac{2\pi}{\lambda} \cdot OPD(z,z_p)} \quad \textbf{(12)}$$

### 2 Strength of reflected & scattered fields

Both electric fields ($\mathbf{E}_{ref}$ and $\mathbf{E}_{sca}$) are related to the illumination field $\mathbf{E}_{illu}$ through the Fresnel reflection $r$, transmission $t$ and the scattering coefficient $s$:

$$\tilde{\mathbf{E}}_{ref}(\mathbf{k}',z=0) = r^{(\perp,\parallel)}(\theta) \cdot \tilde{\mathbf{E}}_{illu}(\mathbf{k}',z=0) \quad \textbf{(13)}$$

$$\tilde{\mathbf{E}}_{sca}(\mathbf{k}',z=0) = s \cdot t_{1,2}^{(\perp,\parallel)}(\theta) \cdot \tilde{\mathbf{E}}_{illu}(\mathbf{k}',z=0) \quad \textbf{(14)}$$

with the subscript $_1$ and $_2$ indicating that the transmission through the glass-water interface has to be accounted for twice. Note that the Fresnel coefficients are defined with respect to $\perp$ & $\parallel$ polarization, i.e. we need to decompose $\mathbf{E}_{illu}$ accordingly:

$$\begin{pmatrix} \tilde{E}_{illu}^{(\perp)}(\mathbf{k}',z=0) \\ \tilde{E}_{illu}^{(\parallel)}(\mathbf{k}',z=0) \\ 0 \end{pmatrix} = \underbrace{\begin{bmatrix} \cos\phi & \sin\phi & 0 \\ -\sin\phi & \cos\phi & 0 \\ 0 & 0 & 1 \end{bmatrix}}_{\hat{\mathbf{P}}(\phi)} \cdot \begin{pmatrix} \tilde{E}_{illu}^{(1)}(\mathbf{k}',z=0) \\ \tilde{E}_{illu}^{(2)}(\mathbf{k}',z=0) \\ 0 \end{pmatrix} \quad \textbf{(15)}$$

Here we have assumed that $\mathbf{E}_{illu}$ is a perfect transverse wave, i.e. fully unpolarized along $z$. In our simulation, $\mathbf{E}_{illu}$ is given as a single plane wave (due to widefield illumination), traveling at $\theta = 0°$ with a certain polarization, e.g. here $x'$-polarized. Hence, we find for the reflection & transmission coefficients, going from glass into water:

$$|r^{(\perp,\parallel)}(\theta)|^2 \rightarrow |r(\theta=0°)|^2 = 4.08\% \quad \textbf{(16)}$$

$$|t_1^{(\perp,\parallel)}(\theta)|^2 \rightarrow |t_1(\theta=0°)|^2 = 99.59\% \quad \textbf{(17)}$$

The scattering coefficient $s$ is related to the scattering cross section $\sigma_{sca}$ through:

$$s = \sqrt{\mu \cdot \frac{\sigma_{sca}}{A}} \quad \textbf{(18)}$$

with $\mu = \arcsin\{\min(NA/n_i, 1)\}$ being the collection efficiency of the objective (*NA* - numerical aperture; $n_i$ refractive index of immersion) [5] and $A$ the area that samples the photon flux, i.e. the effective pixel size of the detector.
The scattering cross section $\sigma_{sca}$, assuming that the protein size is much smaller than $\lambda$, is given as [6]:

$$\sigma_{sca} = \frac{1}{6\pi} \cdot \left(\frac{2\pi}{\lambda} \cdot n_m\right)^4 \cdot |\alpha|^2 \tag{19}$$

with $\lambda$ being the vacuum wavelength of light and $\alpha$ the *polarizability* of the protein. In a first approximation we treat the protein as a spherical particle of homogeneous refractive index $n_p$, for which an analytical expression for $\alpha$ can be found [6]:

$$\alpha = 3 \cdot V_{sphere} \cdot \frac{n_p^2 - n_m^2}{n_p^2 + 2 \cdot n_m^2} \tag{20}$$

with $V_{sphere}$ being the volume of the respective spherical particle with radius $a$:

$$V_{sphere} = \frac{4}{3}\pi \cdot a^3 \tag{21}$$

### 3 Amplitude point-spread functions including near-field components

Next, we compute the amplitude response $\mathbf{h}_{ref}$ by defining the optical systems band limit in Fourier space:

$$\tilde{h}^{(1)}(\mathbf{k}', z=0) = \begin{cases} 1 & |\mathbf{k}'| \leq k_{max} \\ 0 & \text{else} \end{cases} \tag{22}$$

with $k_{max}$ being the maximum transferable spatial frequency given by Abbe's diffraction limit [7] in the fully *coherent* case:

$$k_{max} = \frac{2\pi}{\lambda} \cdot NA \tag{23}$$

By making use of Eq. (6), (8), (11) and $z_p = 0$ (nominal focal plane), we find the corresponding real space representation as:

$$\mathbf{h}_{ref}(\mathbf{r}) = \frac{-i}{\sqrt{\cos\theta}} \cdot \mathcal{F}^{-1}\left\{\hat{\mathbf{T}}(\phi,\theta) \cdot G(z,0) \cdot \tilde{\mathbf{h}}_{ref}(\mathbf{k}', z=0)\right\} \tag{24}$$

With $\sqrt{\cos\theta}$ describing the *aplanatic* factor [2] and $-i$ a phase shift associated with the Huygens-Fresnel principle [8].

Similarly we can compute the amplitude response of the scattering $\mathbf{h}_{sca}$, which can be modeled as the presence of an induced dipole in the protein. Hence, the detectable light is governed by the interaction of such a dipole on top of a refractive index interface. Here we adopt the work of Lieb et al. [9], who describe the electric field distribution of such a dipole at an interface in the *BFP* of the microscope objective. The corresponding fields, in $\perp$ & $\parallel$ polarization, are given as:

$$\begin{aligned} \tilde{h}_{sca}^{(\parallel)}(\mathbf{k}', z=0) &= c_1(\theta) \cdot \sin\theta \cdot \cos\Theta \\ &+ c_2(\theta) \cdot \cos\theta \cdot \sin\Theta \cdot \cos(\phi - \Phi) \end{aligned} \tag{25}$$

$$\tilde{h}_{sca}^{(\perp)}(\mathbf{k}', z=0) = c_3(\theta) \cdot \sin\Theta \cdot \sin(\phi - \Phi) \tag{26}$$

with $\Theta$ and $\Phi$ the azimuth and polar angle of the dipole axis. The constants $c_1 - c_3$ are given according to:

$$c_1(\theta) = n^2 \cdot \frac{\cos\theta}{\cos\theta_s} \cdot t_2^{(\parallel)}(\theta_s) \cdot \Pi(\theta_s) \tag{27}$$

$$c_2(\theta) = n \cdot t_2^{(\parallel)}(\theta_s) \cdot \Pi(\theta_s) \tag{28}$$

$$c_3(\theta) = -n \cdot \frac{\cos\theta}{\cos\theta_s} \cdot t_2^{(\perp)}(\theta_s) \cdot \Pi(\theta_s) \tag{29}$$

$$\Pi(\theta_s) = \exp(i\tfrac{2\pi}{\lambda} n_g \cos\theta_s \delta) \tag{30}$$

where $n = n_m/n_g$ and $\theta_s = \arcsin(n \cdot \sin\theta)$.
Finally, we compute the amplitude response $\mathbf{h}_{sca}$ by setting $z_p = a$ as:

$$\mathbf{h}_{sca}(\mathbf{r}) = \frac{1}{\sqrt{\cos\theta}} \cdot \mathcal{F}^{-1}\left\{\hat{\mathbf{T}}(\phi,\theta) \cdot \hat{\mathbf{P}}(\phi) \cdot G(z,a) \cdot \tilde{\mathbf{h}}_{sca}(\mathbf{k}', z=0)\right\} \tag{31}$$

where we already account for the *Gouy* phase shift [10].

### 4 Information content in the interferometric & ratiometric images

Finally we can express Eq. (1) in terms of the *imaged* electric fields $\mathbf{A}$, i.e. as they appear on the detector:

$$I(\mathbf{r}) = |\mathbf{A}_{ref}|^2 + |\mathbf{A}_{sca}(\mathbf{r})|^2 + 2 \cdot |\mathbf{A}_{ref}| \cdot |\mathbf{A}_{sca}(\mathbf{r})| \cdot \cos\varphi(\mathbf{r}) \tag{32}$$

with $\varphi$ being the phase difference between the scattered and reflected fields at the detector plane, which are given as:

$$\mathbf{A}_{ref} = \mathbf{E}_{ref} \otimes \mathbf{h}_{ref}(\mathbf{r}) \tag{33}$$

$$\mathbf{A}_{sca}(\mathbf{r}) = \mathbf{E}_{sca}(\mathbf{r}) \otimes \mathbf{h}_{sca}(\mathbf{r}) \tag{34}$$

To access the interferometric part of the signal we compute the *ratiometric* contrast $C$, according to [11]:

$$C(\mathbf{r}) = \frac{I(\mathbf{r}) - I_{bkg}}{I_{bkg}} \tag{35}$$

with $I_{bkg}$ being the detectable intensity without the protein:

$$I_{bkg} = |\mathbf{A}_{ref}|^2 \tag{36}$$

Assuming that the scattered field is much weaker than the reflected counterpart, i.e. $|\mathbf{A}_{ref}| \gg |\mathbf{A}_{sca}(\mathbf{r})|$, we get:

$$C(\mathbf{r}) \approx |\mathbf{A}_{sca}(\mathbf{r})| \cdot \frac{2 \cdot \cos\varphi(\mathbf{r})}{|\mathbf{A}_{ref}|} \tag{37}$$

Showing that the detectable contrast is proportional to the scattered field on the detector, which itself is proportional to the polarizability of the protein. The sensitivity of the detection can be increased either by optimizing the phase difference $\varphi$ (such that $\cos\varphi = 1$, i.e. phase matching conditions) or by attenuating the reference field (using a mask in the *BFP*, see [12]).

### 5 Increasing contrast by attenuating & phase shifting the reference field

In principle mass photometry is only limited by shot noise, i.e. the ability to detect a large number of scattered photons. In practice we are limited by the full-well-depth of the detector, which for small proteins is mostly depleted by the reference field. However, as Eq. (37) suggests, we can increase the sensitivity of the detection by decreasing $|\mathbf{A}_{ref}|$. Experimentally this is achieved by placing a mask in the *BFP* of the microscope objective [12]. This is where the scattered and reference fields are separable in terms of the spatial frequency components. We model this by multiplying $\tilde{\mathbf{h}}$ (both: the reference & scattered field) with a Fourier mask $\tilde{\tau}$, given as:

$$\tilde{\tau}(\mathbf{k}') = \begin{cases} |\tilde{\tau}| \cdot e^{i \cdot \arg\{\tilde{\tau}\}} & |\mathbf{k}'| \leq k_{mask} \\ 0 & \text{else} \end{cases} \tag{38}$$

with $|\tilde{\tau}|$ being the transmissivity and $\arg\{\tilde{\tau}\}$ the phase shift of the mask. $k_{mask}$ is related to the spatial extent of the mask with respect of the *BFP* diameter:

$$k_{mask} = \frac{R_{mask}}{f_{obj} \cdot NA} \tag{39}$$

with $R_{mask}$ the radius of the mask in spatial units (e.g. mm).

The results shown in this publication correspond to a mask size that corresponds to an effective *NA* of 0.58. The term *phase matched* refers to an additional phase shift that the mask introduces which was set to $\arg\{\tilde{\tau}\} = \pi/2$, yielding an optimum contrast at the nominal focal plane $z = 0$ (see Fig. 1 b bottom).

### 6 Glass roughness due to phase variations in the reference field

When dealing with real experimental data one typically observes a large, speckle-like, background in the raw data. Following the findings of [13], we modify the constant reference field at the nominal focal plane by a phase variation $\Psi$.

$$I(\mathbf{r}) = \left| \left[ \mathbf{E}_{ref} \cdot e^{i\Psi(\mathbf{r})} \right] \otimes \mathbf{h}_{ref}(\mathbf{r}) + \mathbf{E}_{sca}(\mathbf{r}) \otimes \mathbf{h}_{sca}(\mathbf{r}) \right|^2 \tag{40}$$

This phase distortion is due to the roughness of the glass coverslip. Note that $\Psi$ describes an *effective* phase change, which also accounts for the phase delay of the electric field that transmits through the glass-water interface and eventually leads to the protein scattering. In case of small phase variations, we can approximate the exponential as:

$$e^{i\Psi(\mathbf{r})} \approx 1 + i\Psi(\mathbf{r}) \tag{41}$$

With which we rewrite Eq. (40) into:

$$I(\mathbf{r}) \approx \left|\mathbf{A}_{ref} + \mathbf{A}_{glass}(\mathbf{r}) + \mathbf{A}_{sca}(\mathbf{r})\right|^2 \tag{42}$$

with $\mathbf{A}_{glass}$ being the complex field (at the detector) associated to the glass roughness.

$$\mathbf{A}_{glass}(\mathbf{r}) = \left[i \cdot \mathbf{E}_{ref} \cdot \Psi(\mathbf{r})\right] \otimes \mathbf{h}_{ref}(\mathbf{r}) \tag{43}$$

with $\psi$ being the phase difference between the fields due to glass roughness and protein scattering.
The detected intensity is then given as:

$$\begin{aligned} I(\mathbf{r}) &= |\mathbf{A}_{ref} + \mathbf{A}_{glass}(\mathbf{r}) + \mathbf{A}_{sca}(\mathbf{r})|^2 = \\ &= |\mathbf{A}_{ref}|^2 + |\mathbf{A}_{glass}(\mathbf{r})|^2 + |\mathbf{A}_{sca}(\mathbf{r})|^2 \\ &+ 2 \cdot |\mathbf{A}_{ref}| \cdot |\mathbf{A}_{glass}(\mathbf{r})| \cdot \cos\psi(\mathbf{r}) \\ &+ 2 \cdot |\mathbf{A}_{ref}| \cdot |\mathbf{A}_{sca}(\mathbf{r})| \cdot \cos\varphi(\mathbf{r}) \\ &+ 2 \cdot |\mathbf{A}_{glass}(\mathbf{r})| \cdot |\mathbf{A}_{sca}(\mathbf{r})| \cdot \cos[\psi(\mathbf{r}) - \varphi(\mathbf{r})] \end{aligned} \tag{44}$$

The ratiometric signal Eq. (35) is now given as:

$$C(\mathbf{r}) \approx |\mathbf{A}_{sca}(\mathbf{r})| \cdot 2 \cdot \left[\frac{|\mathbf{A}_{ref}| \cdot \cos\varphi(\mathbf{r})}{|\mathbf{A}_{ref} + \mathbf{A}_{glass}(\mathbf{r})|^2} + \frac{|\mathbf{A}_{glass}(\mathbf{r})| \cdot \cos[\psi(\mathbf{r}) - \varphi(\mathbf{r})]}{|\mathbf{A}_{ref} + \mathbf{A}_{glass}(\mathbf{r})|^2}\right] \tag{45}$$

Alternatively, we could also write the intensity $I$ in terms of an auxiliary reference field $\mathbf{A}'_{ref} = \mathbf{A}_{ref} + \mathbf{A}_{glass}$:

$$I(\mathbf{r}) = |\mathbf{A}'_{ref}(\mathbf{r}) + \mathbf{A}_{sca}(\mathbf{r})|^2 \tag{46}$$

Which we now express with respect to an overall phase shift $\zeta$, comparing the field scattered from the protein with that of the total reference field (including the phase variation due to the glass).

$$\begin{aligned} I(\mathbf{r}) &= |\mathbf{A}'_{ref}(\mathbf{r})|^2 + |\mathbf{A}_{sca}(\mathbf{r})|^2 + 2 \cdot |\mathbf{A}'_{ref}(\mathbf{r})| \cdot |\mathbf{A}_{sca}(\mathbf{r})| \cdot \cos\zeta(\mathbf{r}) = \\ &= |\mathbf{A}_{ref} + \mathbf{A}_{glass}(\mathbf{r})|^2 + |\mathbf{A}_{sca}(\mathbf{r})|^2 + 2 \cdot |\mathbf{A}_{ref} + \mathbf{A}_{glass}(\mathbf{r})| \cdot |\mathbf{A}_{sca}(\mathbf{r})| \cdot \cos\zeta(\mathbf{r}) = \\ &= |\mathbf{A}_{ref}|^2 + |\mathbf{A}_{glass}(\mathbf{r})|^2 + |\mathbf{A}_{sca}(\mathbf{r})|^2 + 2 \cdot |\mathbf{A}_{ref}| \cdot |\mathbf{A}_{glass}(\mathbf{r})| \cdot \cos\psi(\mathbf{r}) \\ &+ 2 \cdot |\mathbf{A}_{ref} + \mathbf{A}_{glass}(\mathbf{r})| \cdot |\mathbf{A}_{sca}(\mathbf{r})| \cdot \cos\zeta(\mathbf{r}) \end{aligned} \tag{47}$$

Which helps us to identify that:

$$\begin{aligned} &|\mathbf{A}_{ref}| \cdot \cos\varphi(\mathbf{r}) + |\mathbf{A}_{glass}(\mathbf{r})| \cdot \cos[\psi(\mathbf{r}) - \varphi(\mathbf{r})] \\ &\quad = |\mathbf{A}_{ref} + \mathbf{A}_{glass}(\mathbf{r})| \cdot \cos\zeta(\mathbf{r}) \end{aligned} \tag{48}$$

Yielding the following for the ratiometric contrast:

$$C(\mathbf{r}) \approx |\mathbf{A}_{sca}(\mathbf{r})| \cdot \frac{2 \cdot \cos\zeta(\mathbf{r})}{|\mathbf{A}_{ref} + \mathbf{A}_{glass}(\mathbf{r})|} \tag{49}$$

Note that this makes the ratiometric contrast vary with the landing position of the particles, which itself leads to mass broadening. A (partial) correction of this can be achieved by computing the following (based on [14]):

$$C'(\mathbf{r}) = C(\mathbf{r}) \cdot \sqrt{I_{bkg}(\mathbf{r})} \tag{50}$$

with the background field given as:

$$I_{bkg}(\mathbf{r}) = \left|\mathbf{A}_{ref} + \mathbf{A}_{glass}(\mathbf{r})\right|^2 \tag{51}$$

This yields the *corrected* ratiometric signal:

$$C'(\mathbf{r}) \approx |\mathbf{A}_{sca}(\mathbf{r})| \quad \cdot \quad 2 \cdot \cos\zeta(\mathbf{r}) \tag{52}$$

This quantity (in principle) still depends on the landing position of the particle, as $\cos\zeta$ is spatially varying due to the glass roughness. Nevertheless, this influence is much weaker (as $|\cos\zeta| \leq 1$). Another practical limitation is the fact that $|\mathbf{A}_{sca}| \propto |\mathbf{E}_{illu}|$, i.e. any inhomogeneous illumination profile will also lead in mass broadening. Hence, proper flat fielding is required, as already discussed in [14].

### 7 Scattering anisotropy of an ellipsoidal particle

When treating *anisotropy* in the polarizability of a protein, we need to reformulate Eq. (19) by replacing the polarizability scalar $\alpha$ with the polarizability tensor $\hat{\alpha}$. In case of shape-dependent anisotropy (e.g. elliptical shape of the protein), the polarizability tensor $\hat{\alpha}$ is defined according to [6]:

$$\hat{\alpha} = \begin{bmatrix} \alpha_a & 0 & 0 \\ 0 & \alpha_b & 0 \\ 0 & 0 & \alpha_c \end{bmatrix} \tag{53}$$

with $\alpha_{a,b,c}$ being the polarizability of an ellipsoidal particle along one of its major axis $a$, $b$ or $c$:

$$\alpha_a = 3 \cdot V_{ellipsoid} \cdot \frac{n_p^2 - n_m^2}{3 \cdot n_m^2 + 3 \cdot L_a \cdot (n_p^2 - n_m^2)} \tag{54}$$

with $V_{ellipsoid}$ being the volume of such an ellipsoid:

$$V_{ellipsoid} = \frac{4}{3}\pi \cdot abc \tag{55}$$

The geometric factors $L_{a,b,c}$ are, e.g. for $a$:

$$L_a = \frac{abc}{2} \cdot \int_0^{+\infty} dv \; \frac{1}{(a^2+v)^{3/2} \cdot (b^2+v)^{1/2} \cdot (c^2+v)^{1/2}} \tag{56}$$

Note that the sum of all three geometric factors yields unity:

$$L_a + L_b + L_c = 1 \tag{57}$$

This means that in the case of a sphere ($a = b = c$), we find that:

$$L_a = L_b = L_c = \frac{1}{3} \tag{58}$$

Yielding exactly the result given in Eq. (20), as then $\alpha_a = \alpha_b = \alpha_c$ and $V_{ellipsoid} \rightarrow V_{sphere}$.
When starting to elongate the sphere on one side (here $a > b = c$), we generate a *prolate* spheroid for which the corresponding geometric factor $L_a$ is given as [6]:

$$L_a = \frac{1-e^2}{e^2} \cdot \left( -1 + \frac{1}{2e} \ln \frac{1+e}{1-e} \right) \tag{59}$$

$$e^2 = 1 - \frac{b^2}{a^2} \tag{60}$$

The corresponding geometric factors $L_b = L_c$ can be found using Eq. (57):

$$L_b = \frac{1}{2} \cdot (1 - L_a) \tag{61}$$

### 8 Deriving the polarizability tensor of a protein from its structure

More interestingly, it is possible to compute the polarizability tensor $\hat{\alpha}$ for a protein, given the knowledge of each atom and its location within the molecule [15, 16] (information which can be obtained from the respective pdb-file of the protein).
The general form of the polarizability tensor $\hat{\alpha}$ is then given by:

$$\hat{\alpha} = \begin{bmatrix} \alpha_{1,1} & \alpha_{1,2} & \alpha_{1,3} \\ \alpha_{2,1} & \alpha_{2,2} & \alpha_{2,3} \\ \alpha_{3,1} & \alpha_{3,2} & \alpha_{3,3} \end{bmatrix} \tag{62}$$

The product of polarizability and electric field equates to a certain, induced, dipole moment $\mathbf{p}$:

$$\mathbf{p}_{exc.} = \underbrace{(\hat{\alpha} - \alpha_m)}_{\text{protein - water}} \cdot \underbrace{t_1^{(\perp,\parallel)}(\theta) \cdot \mathbf{E}_{illu}}_{\mathbf{E}_{sample}} \tag{63}$$

which we have already written as the *excess* dipole moment $\mathbf{p}_{exc.}$, i.e. including the influence of the surrounding medium. Note that $\mathbf{p}$ is a $3 \times 1$ vector, such that:

$$\begin{pmatrix} p^{(1)} \\ p^{(2)} \\ p^{(3)} \end{pmatrix} = \begin{bmatrix} \alpha_{1,1} & \alpha_{1,2} & \alpha_{1,3} \\ \alpha_{2,1} & \alpha_{2,2} & \alpha_{2,3} \\ \alpha_{3,1} & \alpha_{3,2} & \alpha_{3,3} \end{bmatrix} \cdot \begin{pmatrix} E^{(1)}_{sample} \\ E^{(2)}_{sample} \\ E^{(3)}_{sample} \end{pmatrix} \tag{64}$$

The total dipole of such a protein $\mathbf{p}$, is assumed to be a superposition of the individual dipole momenta $\mathbf{p}_i$ of all atoms that make up the molecule:

$$\mathbf{p} = \sum_{i=1}^{N} \mathbf{p}_i(\mathbf{E}_{sample}) \tag{65}$$

According to [16], each of the individual dipole momenta can be expressed as:

$$\mathbf{p}_i(\mathbf{E}_{sample}) = \underbrace{\alpha_i \cdot \mathbf{E}_{sample}}_{\text{atom } i} - \underbrace{\alpha_i \cdot \sum_{j \neq i}^{N} \hat{\mathbf{M}}_{i,j} \cdot \mathbf{p}_j(\mathbf{E}_{sample})}_{\text{all other atoms}} \tag{66}$$

with the dipole field tensor $\hat{\mathbf{M}}$ and $\hat{\mathbf{I}}$ being an identity matrix:

$$\hat{\mathbf{M}}_{i,j} = \begin{cases} \frac{1}{\Delta r^3} \cdot \hat{\mathbf{I}} - \frac{3}{\Delta r^5} \begin{bmatrix} \Delta x^2 & \Delta x \Delta y & \Delta x \Delta z \\ \Delta x \Delta y & \Delta y^2 & \Delta y \Delta z \\ \Delta x \Delta z & \Delta y \Delta z & \Delta z^2 \end{bmatrix} & \Delta r > \rho \\ \frac{4\nu^3 - 3\nu^4}{\Delta r^3} \cdot \hat{\mathbf{I}} - \frac{3\nu^4}{\Delta r^5} \begin{bmatrix} \Delta x^2 & \Delta x \Delta y & \Delta x \Delta z \\ \Delta x \Delta y & \Delta y^2 & \Delta y \Delta z \\ \Delta x \Delta z & \Delta y \Delta z & \Delta z^2 \end{bmatrix} & \text{else} \end{cases} \tag{67}$$

and $\Delta x = x_i - x_j$ (analogous for $\Delta y$ & $\Delta z$), $\Delta r = \sqrt{\Delta x^2 + \Delta y^2 + \Delta z^2}$, $\nu = \Delta r/\rho$, $\rho = 1.662 \cdot (\alpha_i \alpha_j)^{1/6}$.
To further compute the polarizability of the molecule, we can imagine to sequentially illuminate the protein with a plane wave traveling along the $x$, $y$ or $z$-axis, i.e. $\mathbf{E}_l$ with $l \in [x, y, z]$. Yielding a polarizability response in each case as:

$$\hat{\alpha}_l(\mathbf{E}_l) = \begin{pmatrix} \alpha_{l,1} & \alpha_{l,2} & \alpha_{l,3} \end{pmatrix} = \sum_{i=1}^{N} \frac{\mathbf{p}_i(\mathbf{E}_l)}{|\mathbf{E}_l|} \tag{68}$$

We then build up the molecular polarizability tensor $\hat{\alpha}$ from these three responses as:

$$\hat{\alpha} = \begin{bmatrix} \hat{\alpha}(\mathbf{E}_x) \\ \hat{\alpha}(\mathbf{E}_y) \\ \hat{\alpha}(\mathbf{E}_z) \end{bmatrix} = \begin{bmatrix} \alpha_{x,1} & \alpha_{x,2} & \alpha_{x,3} \\ \alpha_{y,1} & \alpha_{y,2} & \alpha_{y,3} \\ \alpha_{z,1} & \alpha_{z,2} & \alpha_{z,3} \end{bmatrix} \tag{69}$$

The work of [15] has shown that it is possible to perform these computations when evaluating large number of atoms (as typical for proteins) when reformulating it in terms of the following matrix equation:

$$\hat{\mathbf{A}} \cdot \hat{\mathbf{p}} = \hat{\mathbf{E}} \tag{70}$$

With $\hat{\mathbf{p}}$ and $\hat{\mathbf{E}}$ being $3N \times 1$ vectors containing the dipole moments and electric fields at each atom ($i \in [1, N]$):

$$\hat{\mathbf{p}} = \begin{pmatrix} \mathbf{p}_1 & \cdots & \mathbf{p}_i & \cdots & \mathbf{p}_N \end{pmatrix}^\top \tag{71}$$

$$\hat{\mathbf{E}} = \begin{pmatrix} \mathbf{E}_1 & \cdots & \mathbf{E}_i & \cdots & \mathbf{E}_N \end{pmatrix}^\top \tag{72}$$

and each individual component given as a $3 \times 1$ vector:

$$\mathbf{p}_i = \begin{pmatrix} p_i^{(1)} & p_i^{(2)} & p_i^{(3)} \end{pmatrix} \tag{73}$$

$$\mathbf{E}_i = \begin{pmatrix} E_i^{(1)} & E_i^{(2)} & E_i^{(3)} \end{pmatrix} \tag{74}$$

The $3N \times 3N$ matrix $\hat{\mathbf{A}}$ is given as:

$$\hat{\mathbf{A}} = \begin{bmatrix} \hat{\alpha}_{1,1}^{-1} & \hat{\mathbf{M}}_{1,2} & \cdots & \hat{\mathbf{M}}_{1,N} \\ \hat{\mathbf{M}}_{2,1} & \hat{\alpha}_{2,2}^{-1} & \cdots & \hat{\mathbf{M}}_{2,N} \\ \vdots & \vdots & \ddots & \vdots \\ \hat{\mathbf{M}}_{N,1} & \hat{\mathbf{M}}_{N,2} & \cdots & \hat{\alpha}_{N,N}^{-1} \end{bmatrix} \tag{75}$$

where each of the elements of $\hat{\mathbf{A}}$ is a $3 \times 3$ matrix, e.g. $\hat{\alpha}^{-1}$ is a diagonal matrix such as:

$$\hat{\alpha}_{i,i} = \begin{bmatrix} \alpha_i^{-1} & 0 & 0 \\ 0 & \alpha_i^{-1} & 0 \\ 0 & 0 & \alpha_i^{-1} \end{bmatrix} \tag{76}$$

In principle, it is now possible to obtain the dipole moments of each atom according to:

$$\hat{\mathbf{p}} = \hat{\mathbf{A}}^{-1} \cdot \hat{\mathbf{E}} \tag{77}$$

With the help of Eq. (68) and (69), we can compute the polarizability tensor $\hat{\alpha}$. In practice this is achieved by iteratively solving Eq. (77) and rewriting $\hat{\mathbf{A}}$ into a sparse matrix through the introduction of a thresholding radius $\Delta r_t$ (here set to 10 Å), which determines the maximum interaction distance between atoms $i$ and $j$ that we take into account [15].

### 9 Mass photometry in the shot-noise limited regime

So far we have only modeled the noise-free signal. Any experiments measuring photon counts are inevitably corrupted by *shot*-noise, which means that the noisy measurement $\mathcal{I}$ is actually given as:

$$\underbrace{\mathcal{I}(\mathbf{r})}_{\substack{\text{noisy}\\ \text{measurement}}} = \underbrace{I(\mathbf{r})}_{\substack{\text{noise-free}\\ \text{expectancy}}} + \underbrace{\mathcal{N}_{\mathcal{I}}(\mathbf{r})}_{\substack{\text{shot}\\ \text{noise}}} \tag{78}$$

with $\mathcal{N}_{\mathcal{I}}$ being the individual noise component in a single measurement and $I$ the expectancy $\langle \mathcal{I} \rangle$:

$$\langle \mathcal{I}(\mathbf{r}) \rangle = I(\mathbf{r}) \tag{79}$$

The noise itself is characterized using a probability distribution connecting the measured outcome $\mathcal{I}$ with the expectancy $I$:

$$P\left[\mathcal{I}(\mathbf{r}) | I(\mathbf{r})\right] = \frac{I(\mathbf{r})^{\mathcal{I}(\mathbf{r})}}{\mathcal{I}(\mathbf{r})!} \cdot e^{-I(\mathbf{r})} \tag{80}$$

which in case of shot noise is given as a Poisson distribution, with variance being equal to the expectancy [17]:

$$\mathit{Var}\{\mathcal{I}(\mathbf{r})\} = \langle \mathcal{I}(\mathbf{r}) \rangle \tag{81}$$

When dealing with large photon counts (as in MP), we can approximate this by a Gaussian distribution:

$$P\left[\mathcal{I}(\mathbf{r}) | I(\mathbf{r})\right] \approx \frac{1}{\sqrt{2\pi \cdot I(\mathbf{r})}} \cdot e^{-\frac{[\mathcal{I}(\mathbf{r}) - I(\mathbf{r})]^2}{2 \cdot I(\mathbf{r})^2}} \tag{82}$$

This shot-noise limitation also translates into a noisy representation of the ratiometric contrast:

$$\underbrace{\mathcal{C}(\mathbf{r})}_{\text{noisy}} = \underbrace{C(\mathbf{r})}_{\text{noise-free}} + \underbrace{\mathcal{N}_{\mathcal{C}}(\mathbf{r})}_{\text{noise}} \tag{83}$$

Assuming that the Gaussian distribution also holds for the ratiometric signal, we write for the probability distribution:

$$P\left[\mathcal{C}(\mathbf{r}) | C(\mathbf{r}); \sigma_{\mathcal{C}}^2(\mathbf{r})\right] \approx \frac{1}{\sqrt{2\pi \cdot \sigma_{\mathcal{C}}^2(\mathbf{r})}} \cdot e^{-\frac{[\mathcal{C}(\mathbf{r}) - C(\mathbf{r})]^2}{2 \cdot \sigma_{\mathcal{C}}^2(\mathbf{r})}} \tag{84}$$

with the measured contrast $\mathcal{C}$, the expectancy $C$ and variance $\sigma_C^2$. Note that in case of ratiometric data the variance is *not* equal to the expectancy anymore, as shown in [18]:

$$\langle \mathcal{C}(\mathbf{r}) \rangle = \frac{\langle \mathcal{I}(\mathbf{r}) \rangle}{\langle \mathcal{I}_{bkg}(\mathbf{r}) \rangle} - 1 = C(\mathbf{r}) \tag{85}$$

$$\sigma_{\mathcal{C}}^2(\mathbf{r}) = Var\{\mathcal{C}(\mathbf{r})\} = \left[\frac{\langle \mathcal{I}(\mathbf{r}) \rangle}{\langle \mathcal{I}_{bkg}(\mathbf{r}) \rangle}\right]^2 \cdot \underbrace{\left[\frac{1}{\langle \mathcal{I}(\mathbf{r}) \rangle} + \frac{1}{\langle \mathcal{I}_{bkg}(\mathbf{r}) \rangle}\right]}_{\approx \frac{2}{\langle \mathcal{I}_{bkg}(\mathbf{r}) \rangle}} \tag{86}$$

with $\langle \mathcal{I}_{bkg}(\mathbf{r}) \rangle$ being the expectancy of the measurement without the protein. Note that we assumed that the noise in the raw intensity data is mainly due to the reflected, not the scattered component (true for small proteins). Substituting $\langle \mathcal{C} \rangle$ yields:

$$Var\{\mathcal{C}(\mathbf{r})\} = [\langle \mathcal{C}(\mathbf{r}) \rangle + 1] \cdot \frac{2}{\langle \mathcal{I}_{bkg}(\mathbf{r}) \rangle} \tag{87}$$

Since $\langle \mathcal{C}(\mathbf{r}) \rangle \ll 1$ we can express the noise variance as:

$$Var\{\mathcal{C}(\mathbf{r})\} \approx \frac{2}{\langle \mathcal{I}_{bkg}(\mathbf{r}) \rangle} \tag{88}$$

Indicating that the noise in the ratiometric distribution is inversely proportional to the expected signal in the raw data $\mathcal{I}$.

### 10 Deriving a basic limit on mass sensitivity & resolution in MP

The further analysis assumes no influence from the glass roughness and a scalar representation of the fields involved, i.e. describes the underlying image formation according to:

$$I(\mathbf{r}) = |A_{ref}|^2 + |A_{sca}(\mathbf{r})|^2 + 2 \cdot |A_{ref}| \cdot |A_{sca}(\mathbf{r})| \tag{89}$$

with $A_{ref}$ and $A_{sca}$ the electric fields on the detector given by:

$$A_{ref} = E_{ref} \otimes h(\mathbf{r}) \tag{90}$$

$$A_{sca}(\mathbf{r}) = E_{sca}(\mathbf{r}) \otimes h(\mathbf{r}) \tag{91}$$

assuming that $h_{ref} = h_{sca} = h$.
We approximate the PSF, corresponding to the intensity signal, as a Gaussian:

$$|h(\mathbf{r})|^2 = h_0^2 \cdot \exp\left(-\frac{|\mathbf{r}|^2}{2 \cdot \sigma_h^2}\right) \tag{92}$$

with $\sigma_h = 0.21 \cdot \lambda / NA$ given in a least-squares sense [19]. The amplitude $h_0^2$ of this Gaussian has the physical meaning of an irradiance (units: $W/m^2$) and can be related to a power $P_0$ (units: $W$) distributed over the *BFP* area $A_{BFP}$ as stated in [20]:

$$h_0^2 = \frac{P_0 \cdot A_{BFP}}{\lambda^2 \cdot f_{obj}^2} \tag{93}$$

The *BFP* area is simply given by integrating a constant field distribution over polar coordinates $\rho$ and $\phi$, with $\rho = f_{obj} \cdot n_i \cdot \sin\theta$.

$$A_{BFP} = \int_0^{\rho_c} d\rho \int_0^{2\pi} d\phi \ \rho = \pi \cdot f_{obj}^2 \quad \cdot \quad \left(n_i \cdot \frac{n_m}{n_g}\right)^2 \tag{94}$$

where $\rho_c$ is the radial distance corresponding to the critical angle $\theta_c = \arcsin(n_m/n_g)$ of the glass - water interface and $f_{obj}$ the focal length of the detection objective, given as the ratio of tube lens focal length to objective magnification: $f_{obj} = f_{tube}/M_{obj}$ . We further modify this *BFP*-area to include the *aplanatic* factor [2] by computing:

$$A_{BFP} = \int_0^{\rho_c} d\rho \int_0^{2\pi} d\phi \ \frac{\rho}{\cos\theta(\rho)} = \int_0^{\rho_c} d\rho \int_0^{2\pi} d\phi \ \frac{\rho}{\cos\left[\arcsin\left(\frac{\rho}{f_{obj} \cdot n_i}\right)\right]} = \pi \cdot f_{obj}^2 \quad \cdot \quad 2 \cdot n_i^2 \cdot \left[1 - \sqrt{1 - \left(\frac{n_m}{n_g}\right)^2}\right] \tag{95}$$

This leads to an effective increase of $A_{BFP}$, which we indicate by the enhancement factor $\gamma$:

$$A_{BFP} \quad \rightarrow \quad \gamma \cdot A_{BFP} \tag{96}$$

with $\gamma = 1.36$ for a 1.42 *NA* oil-immersion objective. Additionally we include the effects of the near-field contribution of the scatterer on the glass-water interface by azimuthally (numerically) averaging Eq. (25) & Eq. (26), comparing the resulting (integrated) field distribution with that of Eq. (94) and find that $\gamma \approx 1.58$ for the same 1.42 *NA* objective.

| *NA* | 1.2 | 1.3 | 1.4 | 1.5 |
|---|---|---|---|---|
| $n_i$ | 1.515 | 1.515 | 1.515 | 1.515 |
| $\gamma$ | 1.14 | 1.21 | 1.53 | 1.67 |

**Table 1.** Enhancement factor $\gamma$ for different detection objective *NA*s.

In terms of the scattering, we write the electric field as a delta-distribution:

$$E_{sca}(\mathbf{r}) = t_2 \cdot t_1 \cdot \delta(\mathbf{r}) \tag{97}$$

The convolution $\delta \otimes h = h$ yields the PSF itself, hence the scattered field on the detector is given as:

$$\begin{aligned} A_{sca}(\mathbf{r}) &= E_{sca}(\mathbf{r}) \otimes h(\mathbf{r}) = \\ &= t_2 \cdot t_1 \cdot \int_{-\infty}^{+\infty} d\mathbf{v} \;\; \delta(\mathbf{v}-\mathbf{r}) \cdot h(\mathbf{v}) = t_2 \cdot t_1 \cdot h(\mathbf{r}) \end{aligned} \tag{98}$$

The scattered power (units $W$) of the protein is given by:

$$P_{sca} = \mu \cdot \sigma_{sca} \cdot I_{illu} \tag{99}$$

with $\mu$ the collection efficiency of the detection objective, $I_{illu}$ the illumination intensity (units: $W/m^2$) and the scattering cross-section (units: $m^2$), given as:

$$\sigma_{sca} = \frac{1}{6\pi} \cdot \left( \frac{2\pi}{\lambda} \cdot n_m \right)^4 \cdot \alpha^2 \tag{100}$$

The polarizability $\alpha$ (units: $m^3$) of a spherical particles is given as:

$$\alpha = 3 \cdot V \cdot \frac{n_p^2 - n_m^2}{n_p^2 + 2 \cdot n_m^2} \tag{101}$$

with $V$ being the volume (units: $m^3$) and $n_p$, $n_m$ the refractive indices of protein and surrounding medium.
Here, however, we want to simply state a linear relationship between the proteins mass $m$ and $\alpha$:

$$\alpha = \delta\alpha \cdot m \tag{102}$$

$$\delta\alpha = 723.857 \, \frac{\text{Å}^3}{\text{kDa}} \tag{103}$$

with $\delta\alpha$ which is the slope of the fitted line in Figure 4a in the main text.
In terms of detected, *scattered*, intensity this yields (note that $t_2$ is already taken care of in $\gamma$):

$$\begin{aligned} |A_{sca}(\mathbf{r})|^2 &= t_1^2 \cdot h_0^2 \cdot \exp\left(-\frac{|\mathbf{r}|^2}{2 \cdot \sigma_h^2}\right) \\ &= \underbrace{t_1^2 \cdot \frac{\mu \cdot \sigma_{sca} \cdot I_{illu} \cdot \gamma \cdot A_{BFP}}{\lambda^2 \cdot f_{obj}^2}}_{|A_{sca}|_0^2} \cdot \exp\left(-\frac{|\mathbf{r}|^2}{2 \cdot \sigma_h^2}\right) \\ &= |A_{sca}|_0^2 \cdot \exp\left(-\frac{|\mathbf{r}|^2}{2 \cdot \sigma_h^2}\right) \end{aligned} \tag{104}$$

Since the illumination is assumed to be widefield, the reflection simply takes the form of a *constant* field distribution:

$$E_{ref} = \tau \cdot r \tag{105}$$

With this we are able to express the detected field $A_{ref}$ as:

$$|A_{ref}|^2 = \underbrace{\tau^2 \cdot r^2 \cdot I_{illu}}_{=|A_{ref}|_0^2} = |A_{ref}|_0^2 \quad \textbf{(106)}$$

Note that both, $A_{ref}$ and $A_{sca}$, are in SI base units of $W/m^2$.
To compute the effect of shot noise, we need to converte the two fields, scattered & reflected, into photon counts. For this we recall the definition of power as the ratio of energy change to time duration required for this change to happen:

$$P = \frac{\Delta E}{\Delta t} \qquad \rightarrow \qquad \Delta E = P \cdot \Delta t \quad \textbf{(107)}$$

In our case $\Delta t$ correspond to the (effective) exposure time when recording light on our detector.
The energy of a single photon at wavelength $\lambda$ is:

$$E_{ph} = h \cdot \frac{c}{\lambda} \quad \textbf{(108)}$$

With this we are able to compute the number of photons per (light) power (units: $W$), according to:

$$N = \frac{\Delta E}{E_{ph}} = \frac{P \cdot \Delta t \cdot \lambda}{h \cdot c} \quad \textbf{(109)}$$

Hence, the number of detected, *reflected*, photons per pixel is now given as:

$$\begin{aligned} N_{ref} &= \mathcal{T} \cdot Q \cdot \left[ \int_{-d_{px}/2}^{+d_{px}/2} dx \int_{-d_{px}/2}^{+d_{px}/2} dy \; |A_{ref}|^2 \right] \cdot \frac{\Delta t \cdot \lambda}{h \cdot c} \\ &= \mathcal{T} \cdot Q \cdot |A_{ref}|_0^2 \cdot \left[ \int_{-d_{px}/2}^{+d_{px}/2} dx \int_{-d_{px}/2}^{+d_{px}/2} dy \; 1 \right] \cdot \frac{\Delta t \cdot \lambda}{h \cdot c} \\ &= \mathcal{T} \cdot Q \cdot |A_{ref}|_0^2 \cdot A_{px} \cdot \frac{\Delta t \cdot \lambda}{h \cdot c} \quad \textbf{(110)} \\ & \quad \textbf{(111)} \end{aligned}$$

with the integration indicating the effect of the photon-sensitive area of a single pixel $A_{px} = d_{px}^2$, $\mathcal{T}$ being the throughput of the optical system and the quantum efficiency $Q$ of the detector.
Similarly we find for the number of detected, *scattered*, photons:

$$\begin{aligned} N_{sca} &= \mathcal{T} \cdot Q \cdot \left[ \int_{-d_{px}/2}^{+d_{px}/2} dx \int_{-d_{px}/2}^{+d_{px}/2} dy \; |A_{sca}|^2 \right] \cdot \frac{\Delta t \cdot \lambda}{h \cdot c} \\ &= \mathcal{T} \cdot Q \cdot |A_{sca}|_0^2 \cdot \left[ \int_{-d_{px}/2}^{+d_{px}/2} dx \int_{-d_{px}/2}^{+d_{px}/2} dy \; \exp\left( -\frac{x^2 + y^2}{2 \cdot \sigma_h^2} \right) \right] \cdot \frac{\Delta t \cdot \lambda}{h \cdot c} \\ &= \mathcal{T} \cdot Q \cdot |A_{sca}|_0^2 \cdot \underbrace{\left[ \sqrt{2\pi} \cdot \sigma_h \cdot \mathrm{erf}\left( \frac{d_{px}}{2\sqrt{2} \cdot \sigma_h} \right) \right]^2}_{\approx A_{px}} \cdot \frac{\Delta t \cdot \lambda}{h \cdot c} \quad \textbf{(112)} \end{aligned}$$

This allows us now to define the photon count with ($N$) and without ($N_{bkg}$) scatterer:

$$\begin{aligned} N &= N_{ref} + N_{sca} + 2 \cdot \sqrt{N_{ref} \cdot N_{sca}} \quad \textbf{(113)} \\ N_{bkg} &= N_{ref} \quad \textbf{(114)} \end{aligned}$$

In the following we are assuming that $N \leq N_{FWD}$, the full-well-depth of the detector. Note that one effectively increases the detected signal beyond $N_{FWD}$ by employing pixel and frame binning (denoted by $n_{bin}$):

$$\begin{aligned} N &= n_{avg} \cdot n_{bin} \cdot \left( N_{ref} + N_{sca} + 2 \cdot \sqrt{N_{ref} \cdot N_{sca}} \right) \quad \textbf{(115)} \\ N_{bkg} &= n_{avg} \cdot n_{bin} \cdot N_{ref} \quad \textbf{(116)} \end{aligned}$$

with $n_{avg}$ being the window size employed in the moving average computation of the ratiometric signal.
The ratiometric contrast in terms of photon counts $C_N$ is:

$$\langle \mathcal{C_N} \rangle = C_N = \frac{N - N_{bkg}}{N_{bkg}} \tag{117}$$

At the same time the variance of this ratiometric signal approximately given as (see Eq. (88)):

$$Var\{\mathcal{C_N}\} \approx \frac{2}{N_{bkg}} \tag{118}$$

Hence, we can define the signal-to-noise ratio *SNR* of the ratiometric signal as:

$$SNR = \frac{\langle \mathcal{C_N} \rangle}{\sqrt{Var\{\mathcal{C_N}\}}} = \frac{1}{\sqrt{2}} \cdot C_N \cdot \sqrt{N_{bkg}} = \frac{1}{\sqrt{2}} \cdot \frac{N - N_{bkg}}{\sqrt{N_{bkg}}} \tag{119}$$

Altogether, the number of detected (*reflected*) photons is given as:

$$N_{bkg} = N_{ref} = I_{illu} \cdot \frac{\Delta t \cdot \lambda}{h \cdot c} \cdot \mathcal{T} \cdot Q \cdot n_{avg} \cdot n_{bin} \cdot \tau^2 \cdot r^2 \cdot d_{px}^2 \tag{120}$$

Similarly, we find for the scattered component ($t_1^2 = 1 - r^2$):

$$N_{sca} = I_{illu} \cdot \frac{\Delta t \cdot \lambda}{h \cdot c} \cdot \underbrace{\frac{(2\pi)^4}{6\pi}}_{=\frac{8}{3}\pi^3} \cdot n_{avg} \cdot n_{bin} \cdot \frac{\mathcal{T} \cdot Q \cdot \left[1 - r^2\right] \cdot \mu \cdot n_m^4 \cdot \gamma \cdot A_{BFP} \cdot d_{px}^2}{\lambda^6 \cdot f_{obj}^2} \cdot [\delta\alpha \cdot m]^2 \tag{121}$$

The interference component is now defined as:

$$2 \cdot \sqrt{N_{ref} \cdot N_{sca}} = I_{illu} \cdot \frac{\Delta t \cdot \lambda}{h \cdot c} \cdot 2\sqrt{\frac{8}{3}\pi^3} \cdot n_{avg} \cdot n_{bin} \cdot \frac{\mathcal{T} \cdot Q \cdot \sqrt{1 - r^2} \cdot \sqrt{\mu} \cdot n_m^2 \cdot \sqrt{\gamma} \cdot \sqrt{A_{BFP}} \cdot d_{px}^2 \cdot \tau \cdot r}{\lambda^3 \cdot f_{obj}} \cdot \delta\alpha \cdot m \tag{122}$$

Hence, the differential signal, neglecting the purely scattering component, is given as:

$$\begin{aligned} N - N_{bkg} &\approx 2 \cdot \sqrt{N_{ref} \cdot N_{sca}} = \\ &= I_{illu} \cdot \frac{\Delta t \cdot \lambda}{h \cdot c} \cdot 2\sqrt{\frac{8}{3}\pi^3} \cdot n_{avg} \cdot n_{bin} \cdot \frac{\mathcal{T} \cdot Q \cdot \sqrt{1 - r^2} \cdot \sqrt{\mu} \cdot n_m^2 \cdot \sqrt{\gamma} \cdot \sqrt{A_{BFP}} \cdot d_{px}^2 \cdot \tau \cdot r}{\lambda^3 \cdot f_{obj}} \cdot \delta\alpha \cdot m \end{aligned} \tag{123}$$

And the square-root of the reflected component as:

$$\sqrt{N_{bkg}} = \sqrt{I_{illu}} \cdot \sqrt{\frac{\Delta t \cdot \lambda}{h \cdot c}} \cdot \sqrt{n_{avg} \cdot n_{bin}} \cdot \sqrt{\mathcal{T} \cdot Q} \cdot \tau \cdot r \cdot d_{px} \tag{124}$$

Finally we are able to write for the ratiometric SNR:

$$\begin{aligned} SNR &= \frac{1}{\sqrt{2}} \cdot \frac{N - N_{bkg}}{\sqrt{N_{bkg}}} \approx \frac{2}{\sqrt{2}} \cdot \sqrt{N_{sca}} = \\ &= \sqrt{I_{illu}} \cdot \sqrt{\frac{\Delta t \cdot \lambda}{h \cdot c}} \cdot \frac{2}{\sqrt{2}} \sqrt{\frac{8}{3}\pi^3} \cdot \sqrt{n_{avg} \cdot n_{bin}} \cdot \frac{\sqrt{\mathcal{T} \cdot Q} \cdot \sqrt{1 - r^2} \cdot \sqrt{\mu} \cdot n_m^2 \cdot \sqrt{\gamma} \cdot \sqrt{A_{BFP}} \cdot d_{px}}{\lambda^3 \cdot f_{obj}} \cdot \delta\alpha \cdot m \end{aligned} \tag{125}$$

Note that the *SNR* is independent of the mask strength $\tau$, as it influences contrast and noise simultaneously, such that they cancel each other. Contrary to this, the ratiometric contrast is improved upon modifying the effective reflectivity from the glass-buffer medium interface:

$$\langle \mathcal{C_N} \rangle = 2\sqrt{\frac{8}{3}\pi^3} \cdot \frac{\sqrt{1 - r^2} \cdot \sqrt{\mu} \cdot n_m^2 \cdot \sqrt{\gamma} \cdot \sqrt{A_{BFP}}}{\lambda^3 \cdot f_{obj} \cdot \tau \cdot r} \cdot \delta\alpha \cdot m \tag{126}$$

But the contrast is not simply enhanced by increasing $I_{illu}$, $n_{avg}$, $n_{bin}$ or any other parameter that yields a higher photon flux at the detector. Those merely affect the noise of the ratiometric signal, as:

$$Var\{\mathcal{C_N}\} \approx \frac{2}{I_{illu} \cdot \frac{\Delta t \cdot \lambda}{h \cdot c} \cdot \mathcal{T} \cdot Q \cdot n_{avg} \cdot n_{bin} \cdot \tau^2 \cdot r^2 \cdot d_{px}^2} \tag{127}$$

We can use the analytical model to compute a *mass-equivalent* signal-to-noise ratio: the smallest mass $m_q$ that one would be able to detect with an SNR-level of $q$:

$$m_q = \frac{q}{\delta\alpha} \cdot \frac{1}{\sqrt{I_{illu}}} \cdot \sqrt{\frac{h \cdot c}{\Delta t \cdot \lambda}} \cdot \frac{\sqrt{2}}{2} \sqrt{\frac{3}{8\pi^3}} \cdot \frac{1}{\sqrt{n_{avg} \cdot n_{bin}}} \cdot \frac{\lambda^3 \cdot f_{obj}}{\sqrt{\mathcal{T} \cdot Q} \cdot \sqrt{1-r^2} \cdot \sqrt{\mu} \cdot n_m^2 \cdot \sqrt{\gamma} \cdot \sqrt{A_{BFP}} \cdot d_{px}} \tag{128}$$

In terms of mass *resolution* we employ the concept of the Quantum-Cramer-Rao lower bound (QCRLB) [21]. The smallest achievable uncertainty of estimating the mass of the protein is then given as:

$$\begin{aligned} \sigma_m &= \frac{1}{2} \cdot \frac{m}{\sqrt{N_{sca}}} = \frac{m}{\sqrt{2}} \cdot \frac{1}{SNR} = m_{q=\sqrt{0.5}} = \qquad (129) \\ &= \frac{1}{2} \cdot \frac{1}{\delta\alpha} \cdot \frac{1}{\sqrt{I_{illu}}} \cdot \sqrt{\frac{h \cdot c}{\Delta t \cdot \lambda}} \cdot \sqrt{\frac{3}{8\pi^3}} \cdot \frac{1}{\sqrt{n_{avg} \cdot n_{bin}}} \cdot \frac{\lambda^3 \cdot f_{obj}}{\sqrt{\mathcal{T} \cdot Q} \cdot \sqrt{1-r^2]} \cdot \sqrt{\mu} \cdot n_m^2 \cdot \sqrt{\gamma} \cdot \sqrt{A_{BFP}} \cdot d_{px}} \end{aligned}$$

Note that this uncertainty independent on whether the measurement is performed interferometrically or in darkfield mode. The latter, however, exhibits a worsened SNR by a factor of $\sqrt{2}$, as:

$$SNR_{darkfield} = \frac{N_{sca}}{\sqrt{N_{sca}}} = \sqrt{N_{sca}} = \frac{1}{\sqrt{2}} \cdot SNR \tag{130}$$

with:

$$\sqrt{N_{sca}} = \sqrt{I_{illu}} \cdot \sqrt{\frac{\Delta t \cdot \lambda}{h \cdot c}} \cdot \sqrt{\tfrac{8}{3}\pi^3} \cdot \sqrt{n_{avg} \cdot n_{bin}} \cdot \frac{\sqrt{\mathcal{T} \cdot Q} \cdot \sqrt{1-r^2} \cdot \sqrt{\mu} \cdot n_m^2 \cdot \sqrt{\gamma} \cdot \sqrt{A_{BFP}} \cdot d_{px}}{\lambda^3 \cdot f_{obj}} \cdot \delta\alpha \cdot m \tag{131}$$

Making it more difficult to reach the fundamental $\sigma_m$-value in practice. Of course this also translates into the smallest detectable mass: $m_{q,darkfield} = \sqrt{2} \cdot m_q$.

### 11 Signal-to-noise ratio including excess noise:

To including the additional excess (or baseline) noise we add the additional uncertainty $\sigma^2_{exc.}$ (in terms of the ratiometric signal) to the variance stemming from shot-noise, according to:

$$SNR_{exc.} = \frac{\langle \mathcal{C}_\mathcal{N} \rangle}{\sqrt{Var\{\mathcal{C}_\mathcal{N}\} + \sigma^2_{exc.}}} = \frac{C_N}{\sqrt{\frac{2}{N_{bkg}} + \sigma^2_{exc.}}} \tag{132}$$

We can rearrange the denominator which yields in:

$$\sqrt{\frac{2}{N_{bkg}} + \sigma^2_{exc.}} = \sqrt{\frac{2 + \sigma^2_{exc.} \cdot N_{bkg}}{N_{bkg}}} \tag{133}$$

Altogether we find for the signal-to-noise ratio including the excess noise:

$$\begin{aligned} SNR_{exc.} &= \frac{N - N_{bkg}}{N_{bkg}} \cdot \sqrt{N_{bkg}} \cdot \frac{1}{\sqrt{2 + \sigma^2_{exc.} \cdot N_{bkg}}} = \qquad (134) \\ &= \frac{N - N_{bkg}}{\sqrt{N_{bkg}}} \cdot \frac{1}{\sqrt{2 + \sigma^2_{exc.} \cdot N_{bkg}}} \qquad (135) \end{aligned}$$

Which we can relate to the shot-noise limited *SNR* through:

$$SNR_{exc.} = SNR \cdot \underbrace{\frac{\sqrt{2}}{\sqrt{2 + \sigma^2_{exc.} \cdot N_{bkg}}}}_{=\xi} \tag{136}$$

Verifying that for $\sigma_{exc.} = 0$ we get: $SNR_{exc.} = SNR$. Note that this automatically means that the $SNR_{exc.}$ equivalent mass $m_{q,exc.}$ is still given by Eq. (128), only with the substitution:

$$q \quad \rightarrow \quad q/\xi \tag{137}$$

Finally we would want to see how $SNR_{exc.}$ behaves when detecting a larger number of photons, e.g. through increase of the illumination (in case of pure shot-noise the SNR is unlimited as: $SNR^{max} = +\infty$). For this we introduce a scaling factor $\epsilon$:

$$N \rightarrow \epsilon \cdot \left( N_{bkg} + N_{sca} + 2 \cdot \sqrt{N_{bkg} \cdot N_{sca}} \right) \quad \textbf{(138)}$$

$$N_{bkg} \rightarrow \epsilon \cdot N_{bkg} \quad \textbf{(139)}$$

So we get for the shot-noise limited $SNR$:

$$\sqrt{2} \cdot SNR \approx \frac{\epsilon \cdot 2 \cdot \sqrt{N_{bkg} \cdot N_{sca}}}{\sqrt{\epsilon \cdot N_{bkg}}} = \sqrt{\epsilon} \cdot 2 \cdot \sqrt{N_{sca}} \quad \textbf{(140)}$$

With this we find for $SNR_{exc.}$:

$$SNR_{exc.} = 2 \cdot \sqrt{\frac{\epsilon \cdot N_{sca}}{2 + \sigma_{exc.}^2 \cdot \epsilon \cdot N_{bkg}}} = 2 \cdot \sqrt{\frac{N_{sca}}{\frac{2}{\epsilon} + \sigma_{exc.}^2 \cdot N_{bkg}}} \quad \textbf{(141)}$$

Taking this in the limit of $\epsilon \rightarrow \infty$:

$$\lim_{\epsilon \rightarrow \infty} SNR_{exc.} = \frac{2}{\sigma_{exc.}} \cdot \sqrt{\frac{N_{sca}}{N_{bkg}}} \quad \textbf{(142)}$$

With the following ratio of scattered to reflected number of photons:

$$\frac{N_{sca}}{N_{ref}} = \frac{|A_{sca}|_0^2}{|A_{ref}|_0^2} = \frac{t_1^2 \cdot \frac{\mu \cdot \sigma_{sca} \cdot \gamma \cdot A_{BFP}}{\lambda^2 \cdot f_{obj}^2}}{\tau^2 \cdot r^2} =$$

$$= \frac{t_1^2 \cdot \mu \cdot (2\pi)^4 \cdot n_m^4 \cdot \gamma \cdot A_{BFP}}{\lambda^6 \cdot f_{obj}^2 \cdot \tau^2 \cdot r^2} \quad \textbf{(143)}$$

### 12 Mass photometry signal for different mask types

Figure 1 presents line profiles of the simulated data for Hsp16.5 (24-mer; $\approx 400$ kDa) in the lateral (a & c) and axial (b) dimensions. In general increasing the mask strength increases the sensitivity of the MP-system as the small ratiometric contrast (a & b) gets enhanced, such that the signal rises above the shot-noise level. To obtain the optimum performance it is necessary to further make sure that the reference and scattered light are exactly in-phase, which requires the mask to introduce an additional $\pi/2$ phase shift. We term this the *phase matched* mask (a & b, bottom) and now observe the optimum contrast and the nominal focal plane. Note that, when considering a finite full-well-depth of the detector, the attenuation mask can not be arbitrarily strong. As the recorded modulation due to glass roughness increases such, that certain pixels will be saturated.

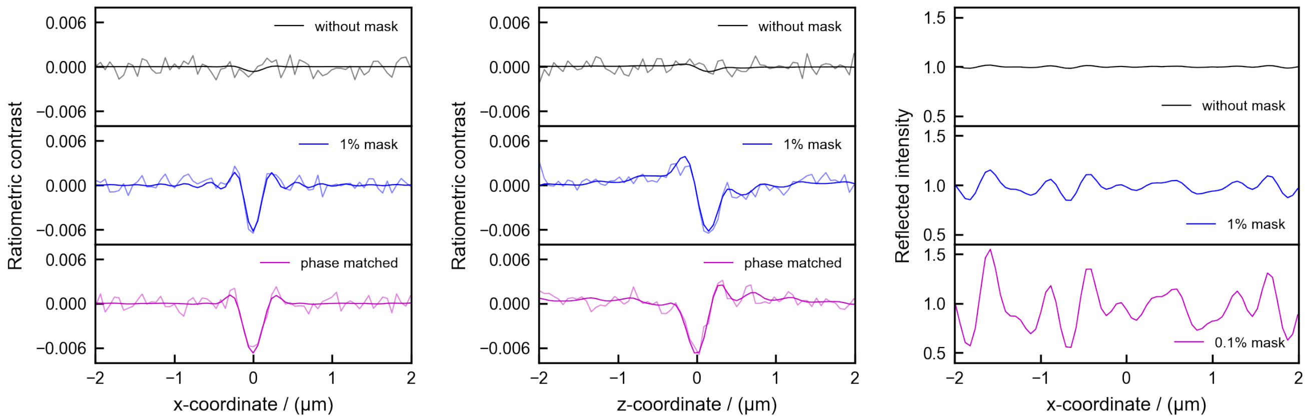


**(a)** **(b)** **(c)**

**Fig. 1.** Ratiometric contrast and recorded intensity when simulating the MP signal of Hsp16.5 (24-mer, $\approx 400$ kDa). a) Lateral line profile of the ratiometric contrast, when imaging using no mask (top, black), a $1\%$ mask (middle, blue) and a $1\%$ mask with phase matching condition (bottom, magenta). The thick line represents the expected (noise-free) signal and the thin line a noisy measurement. (b) The same as in (a), only as a axial line profile. Note how the axial position of optimum contrast is shifted towards the nominal focal plane, when using the phase matched $1\%$ mask. (c) Lateral line profile of the recorded intensity, including the effects due to glass roughness. Improving the performance of the MP-system by increasing the mask strength is limited, as this leads to a stronger detected modulation in the raw data, which ultimately fills the complete full-well-depth of the detector, i.e. preventing more information to be captured.

### 13 Comparison between PDB vs alphafold structure of BSA

As noted in the main text, the pdb-file that we used to simulate the ratiometric signal of BSA (PDBID 3V03) does only contain a smaller subset of the atoms of a real BSA molecule in buffer solution. To get a more realistic estimation of the inferred contrast we performed the same simulation with the polarizability obtained from a pdb-structured predicted by alphafold [22] (UniProt P02769). Overall we observe an increase of $\approx 11\%$, resulting in $m \approx 64$ kDa, in good agreement with the experimentally expected values.

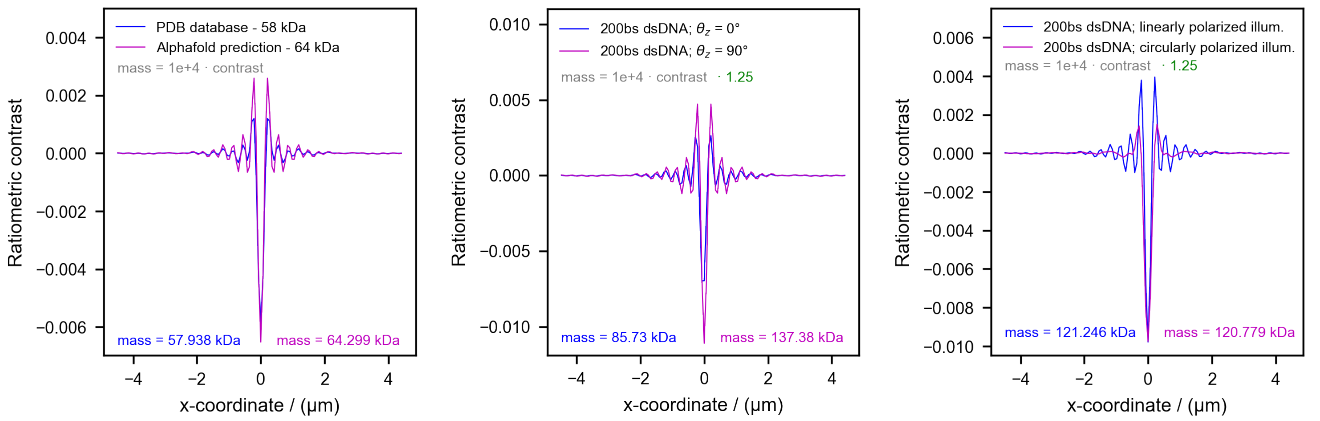


**(a)** **(b)** **(c)**

**Fig. 2.** Ratiometric contrast comparison between pdb & alphafold prediction of BSA (a), 200 bp double-stranded (ds) DNA at different orientations (b) and illumination polarizations. a) The alphafold prediction of BSA (UniProt P02769) contains the atoms that were missing in the original pdb structure (PDBID 3V03), yielding an overall increase of the inferred mass by $\approx 11\%$. b) When imaging 200bp dsDNA using linearly polarized illumination the ratiometric signal changes with the orientation ($\theta_z = 0^\circ$, $\theta_z = 90^\circ$) of the rod-like molecule. c) Ratiometric signal for randomly oriented dsDNA molecule imaged with linearly (blue) or circularly (magenta) polarized light, yielding the same contrast which corresponds to $\approx 121$ kDa. Note that the mass-to-contrast conversion is larger when imaging dsDNA, compared to proteins (see [23]).

### 14 MP signal for 200bp dsDNA imaged with linearly & circularly polarized illumination

In Figure 3 a of the main text we show experimental results of imaging 200 base pair (bp) double-stranded (ds) DNA. We further investigated the theoretical signal to be expected from such a rod-like structure. For this we first generated a random sequence of dsDNA containing 200 bp which we transferred into a pdb-file (B-DNA model) using [24]. The simulated ratiometric signal for two orthogonal orientations ($\theta_z = 0^\circ$, $\theta_z = 90^\circ$) are shown in Fig. 2 b), indicating a strong change depending how the rod-like molecule is aligned with the linearly polarized illumination. We also investigate the effect of randomly oriented dsDNA (200 bp) when being imaged either with linearly (blue) or circularly (magenta) polarized light. Both imaging scenarios yield the same ratiometric contrast amounting to $\approx 121$ kDa. Note the larger mass-to-contrast conversion factor ($\times 1.25$) when imaging dsDNA compared to that of proteins, as reported in [23].

### 15 Simulated mass resolution for detecting BSA monomer at different exposure times

Being able to simulate landing assay movies we looked into the mass resolution, i.e. the width of the fitted Gaussian, when detecting the BSA-monomer alone. We do this for a changing effective exposure time and assume shot-noise limited performance. The results are shown in Fig. 3 a) for different simulated scenarios and experimental results.
Starting out with a $0.1\%$ mask and simulated glass roughness without the correction in post-processing (black curve) we observe an optimum value of $\sigma_m \approx 5.5$ kDa. Lowering the effective exposure time will make the analyzed ratiometric movies look more noisy, making it more difficult for the particle picking & fitting algorithms to perform optimally. Making use of the correction step (Eq. (50); blue) enables to extract a minimum $\sigma_m \approx 2$ kDa, in this ideal simulated scenario. Interestingly, performing the simulation without the glass roughness (hence no need for such a correction; green), yields a similar value for $\sigma_m$. Indicating that in principle the correction is enabling to achieve a performance similar to that of using a perfectly flat coverslip, assuming a constant $|\mathbf{E}_{illu}|$. Experimental results are shown in red, exhibiting a minimum $\sigma_m \approx 7.5$ kDa, which fits more to a $1\%$ mask. This difference is most likely due to the additional baseline noise of $\approx 5$ kDa (see Fig. 5 in main text).

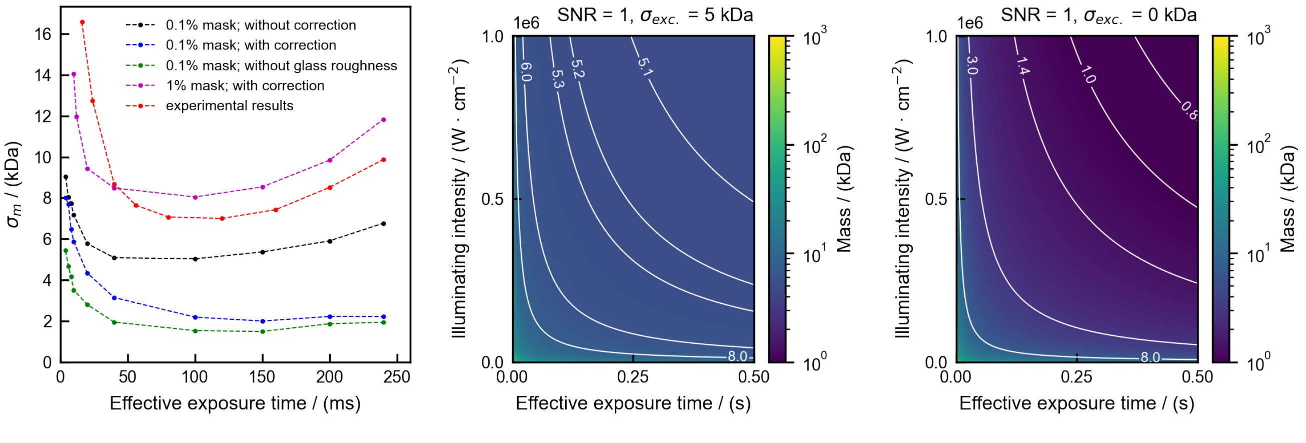


**Fig. 3.** a) Fitted width of Gaussian for simulated and experimental landing assay data of the BSA monomer, in dependency of effective exposure time. Simulating a $0.1\%$ mask including glass roughness yields a minimum $\sigma \approx 5.5$ kDa, while this can be further decreased when making use of the correction step in post-processing to $\sigma \approx 2$ kDa. Note that this comes very close to the value we observe when simulating without glass roughness. Experimental results, however, achieve $\sigma \approx 7.5$ kDa, which most likely is due to the additional baseline noise of $\approx 5$ kDa as described in the main text (e.g. see Fig. 5 b - c) Minimum detectable mass with $SNR = 1$ with (b) and without (c) this additional excess noise. For realistic imaging parameters we find that $m_{q=1} \approx 5$ kDa, which is in agreement with the results shown in [25].

### 16 Smallest detectable mass with $SNR = 1$; with and without excess noise

Equation Eq. (128) enables us to calculate the smallest detectable mass given a certain $SNR$ level. Here we show the results for the fundamental limit, i.e. $SNR = 1$. When including the additional baseline noise of $\approx 5$ kDa and realistic imaging parameters, we observe $m_{q=1} \approx 5$ kDa. Which seems realistic given the results shown in [25], detecting a $9$ kDa protein at $SNR \approx 1.4$. When neglecting the additional baseline noise our calculation suggest to reach $m_{q=1} \approx 1.5$ kDa with the same illumination power & effective integration time.
